# A mathematical model of the HPA axis demonstrates bifurcations under varying stress and systemic conditions

**DOI:** 10.64898/2026.10.07.757338

**Authors:** I. G. Violaris, D. Pinotsis, M. Tsipouras, K. Kalafatakis

## Abstract

In this study, we develop a mathematical model of the hypothalamic–pituitary–adrenal (HPA) axis and systematically investigate the model complexity required to reproduce observed hormone dynamics. The model is fitted to ACTH and cortisol measurements from five healthy volunteers reported in the literature. We show that simultaneous reproduction of both hormone profiles requires a delay differential equation model with at least five state variables among the model architectures considered. Following fitting the volunteer data, we investigate the response of the model to sustained stress using bifurcation analysis. The coupled system exhibits bifurcations over a range of stress intensities. This analysis reveals bifurcations whose critical stress thresholds depend strongly on the modulatory state of the system. These results provide a mathematical framework for investigating transitions between physiological and dysregulated HPA axis dynamics, which may be related to states of health and disease.

## Introduction

The dynamics of cortisol have been studied in both rats and humans since the early twentieth century [1] [2]. Cortisol appears to exhibit both ultradian rhythms (with a period of less than 24 hours) and circadian rhythms (with a period of approximately twenty-four hours) [3], both of which constitute important determinants of its biological action. The proper function of the hypothalamic–pituitary–adrenal (HPA) axis, which is responsible for cortisol production in the body, is necessary for a variety of cellular homeostatic functions [4] and for the optimal adaptation of the organism to stressful conditions [5]. Mathematical modelling research conducted to date has focused primarily on systems of ordinary differential equations and delay differential equations attempting to describe hormonal dynamics under both physiological and pathological conditions, including Addison’s disease, adrenal insufficiency, hypercortisolemia, hypocortisolemia, and related disorders[6], [7], [8] [9] .

The conventional textbook representation of cortisol regulation is typically presented as a relatively simple feedback loop [10]. Corticotropin-releasing hormone (CRH) is released from the hypothalamus in response to signals received by the paraventricular nucleus (PVN), stimulating the pituitary gland to secrete adrenocorticotropic hormone (ACTH). ACTH then enters the circulation and, after a delay of approximately 20 minutes, stimulates cortisol production by the adrenal glands. Circulating cortisol subsequently exerts negative feedback on ACTH secretion, thereby closing the regulatory loop [11]. This schematic description, however, represents only the coarse structure of the HPA axis. The biochemical mechanisms governing glucocorticoid feedback and adrenal responsiveness are considerably more complex and involve multiple intermediate regulatory processes whose activity varies over time. These mechanisms motivated the introduction of the time-dependent modulatory functions *E*_1_(*t*) and *E*_2_(*t*) in our model. The principal biological processes underlying these terms are described in greater detail in the following section.

Cortisol binds the glucocorticoid receptor; following nuclear translocation, activated GR can transrepress Nur77/Nurr1-mediated activation of POMC transcription, thereby reducing ACTH synthesis [10] [12]. Schematically, this may be represented as follows: This process is modulated by clock gene activity as shown in Charmandari et al [13]. Thus the naive approach where constants are used is not enough to properly encapsulate the process. Following this, steroidogenesis is also an extremely complex process relying on cyclic AMP (cAMP) and steroidogenic acute regulatory (StAR) protein availability. The following schematic based on Miller and Auchus [14] shows a simplified account of how this process unfolds.

The multiscale nature of these regulatory mechanisms, together with the presence of interacting feedback processes and physiological delays, motivates a dynamical description of the HPA axis. The physiological state of the HPA axis cannot be adequately characterised by static measures of cortisol concentration alone. Cortisol secretion exhibits pronounced ultradian pulsatility superimposed upon circadian variation, while its temporal profile emerges from feedback interactions between the components of the HPA axis, physiological delays, and time-dependent modulation of both feedback and adrenal responsiveness. Consequently, similar average cortisol concentrations may correspond to markedly different underlying dynamical states. Inter-individual variation in these temporal patterns, together with the transient response of the axis to stress, further motivates treating HPA regulation as a dynamical-systems problem rather than solely as a problem of predicting hormone concentrations at equilibrium or through summary statistics.

Thus, in this study we develop a mathematical model of the HPA axis that explicitly accounts for its dynamical complexity. We begin with a minimal model and show that its architecture is insufficient to reproduce the observed hormone dynamics. We then extend the model to a structure capable of capturing the temporal behaviour of the axis and further refine it through the introduction of two time-dependent modulatory functions. These functions are constrained so that their waveforms do not simply reproduce or track the measured hormone profiles. Instead, they serve two related purposes. Firstly, they provide a phenomenological representation of time-varying regulatory processes not explicitly included in the core model. Secondly, they provide modulatory variables through which changes in the regulatory state of the system can be investigated, including their effects on its bifurcation structure. A fully mechanistic biological formulation of these regulatory processes is non-trivial and lies beyond the scope of the present study. After obtaining successful fits for five healthy volunteers, we investigate the resulting model dynamics and their physiological implications. This analysis reveals bifurcations across varying systemic and stress conditions which appear when there exists a mismatch between modulating functions and the core HPA axis model.

## Results

We developed three successive models, the results of which are presented in this section. The successive model formulations considered below are not introduced primarily to optimize goodness of fit, but to progressively address structural and physiological limitations of the preceding formulations and thereby obtain a dynamical representation suitable for mechanistically interpretable bifurcation analysis. Improved agreement with the experimental measurements emerges as a consequence of this model-development process. The purpose of model calibration was therefore not to optimize agreement with the data as an end in itself, but to constrain successive model architectures against human observations so that their subsequent dynamical analysis could be interpreted in relation to HPA-axis physiology.

### Minimal model

The minimal model represents the HPA axis using only its three principal hormonal variables: CRH, ACTH, and cortisol. It constitutes the simplest formulation considered here in which CRH is represented explicitly as a dynamical variable rather than prescribed directly as an external input. This model is used to determine how much of the observed HPA-axis dynamics can be reproduced by the core hormonal interactions alone.

We present only a single simulation of the minimal model, since this formulation serves primarily as an intermediate step toward the more complete model architectures considered below. All fits in this study use data from Upton et al. [18]. For the minimal model, we considered Healthy Volunteer 1 only. In the subsequent mechanistic and modulated models, we fitted data from five volunteers.

Upton et al. reported hormone profiles for seven volunteers; however, Volunteers 3 and 7 were excluded from the present analysis because their datasets contained substantial missing data or apparent measurement artefacts. Despite its simplicity, the minimal model is able to reproduce the overall structure of the observed hormone profiles to a reasonable extent.

As illustrated by this simulation, the minimal model is capable of generating oscillatory dynamics and reproducing the broad structure of the observed hormone profiles. However, this agreement is achieved only at the cost of unrealistically large temporal lags between the ACTH and cortisol peaks. This limitation is not a consequence of the fitting procedure, but follows directly from the mathematical structure of the model (see Supplementary Appendix A). In particular, within this model architecture, the minimal feedback architecture cannot generate autonomous ultradian Hopf oscillations having both the observed period and the physiological ACTH-cortisol lag. A more complex dynamical architecture is therefore required (see Supplementary Appendix A).

### Mechanistic model

The mechanistic model extends the minimal formulation by introducing two intermediate glucocorticoid-receptor states, *G*_1_ and *G*_2_, between cortisol and the inhibition of ACTH production. These additional states provide a more explicit representation of glucocorticoid-mediated negative feedback and distribute the temporal response across intermediate regulatory processes, addressing the unrealistically large ACTH-to-cortisol lag required by the minimal model. Studies of human physiology indicate that circulating ACTH stimulates adrenal cortisol secretion after a delay of approximately 10–20 minutes [**?**]. In the hormone profiles modelled here [**?**], the observed ACTH–cortisol lag was approximately 20 minutes.

The five-equation mechanistic model produces substantially better fits than the minimal model, although there remains clear scope for further improvement. In general, we preferred fits in which the model underestimated some of the observed peaks rather than introducing additional unmeasured forcing terms.

The CRH signal, *R*, is driven here only by the light-related input, whereas in reality it may also be influenced by factors such as central nervous system stress, feeding, and other physiological signals considered later in the exploratory analysis. Because no direct CRH measurements were available for these volunteers, attempting to infer such additional inputs would lead to a severely underdetermined problem. Introducing sufficiently flexible latent CRH forcing could substantially improve the fit to cortisol, but at the cost of poor identifiability and reduced predictive interpretability.

We therefore chose not to introduce additional unconstrained CRH inputs. Representative fits for Volunteers 1 and 2 are presented here, while the remaining fits are provided in Supplementary Appendix C .

As shown by these simulations, the five-equation mechanistic model reproduces the observed hormone profiles substantially better than the minimal model. In addition, the resulting ACTH– cortisol lag is physiologically appropriate in most cases i.e. within the observed 10 to 20 minute lag. The remaining volunteer fits are presented in Supplementary Appendix D.

### Modulated mechanistic model

The Fourier-modulated model retains the mechanistic core of the five-state formulation while introducing the time-dependent modulatory functions *E*_1_(*t*) and *E*_2_(*t*). These functions provide phenomenological representations of time-varying regulatory processes that are not explicitly captured by the core model architecture, allowing their influence on the observed hormonal dynamics and on the dynamical behaviour of the system to be investigated.

Finally, we present the results of the full model incorporating the two Fourier-based modulatory functions. This formulation provides a substantially improved fit to the data compared with the two preceding model architectures. The increased flexibility, however, comes at a cost in identifiability, since the inferred modulatory signals may represent the combined effects of several unmeasured processes, including clock-gene-dependent regulation, latent CRH inputs, measurement noise, cAMP dynamics, and StAR availability.

The Fourier functions were therefore deliberately kept simple and constrained during fitting. Their purpose was not to track the measured hormone profiles directly, but to capture only the minimum additional time-dependent modulation required to improve the fit while limiting the risk of overfitting.

The remaining simulations are presented in Supplementary Appendix C. The inclusion of the modulatory functions *E*_1_(*t*) and *E*_2_(*t*) improves the fits substantially. For example, in Volunteer 2 the purely mechanistic model generates two pulses where the experimental profile contains only one, whereas the modulated model reproduces the observed pattern more accurately. The inferred modulatory functions for this volunteer exhibit a comparatively complex waveform, but their behaviour remains compatible with the type of smooth oscillatory signal that could plausibly arise from an underlying biological oscillator.

### Stress response and qualitative changes in model dynamics

The purpose of this experiment was to determine whether the fitted system exhibited homeostatic recovery following a transient disturbance, defined here as a return towards the corresponding unstressed trajectory after removal of the perturbation, when evaluated using the model parameter set fitted to the Healthy Volunteer 1 data.

As shown in Fig. 8, the perturbation produced a transient displacement of the ACTH and cortisol trajectories, after which the system returned towards the corresponding unstressed trajectory. Thus, for the fitted Healthy Volunteer 1 parameterization, the model exhibited homeostatic recovery following an acute perturbation.

**Figure 1.**
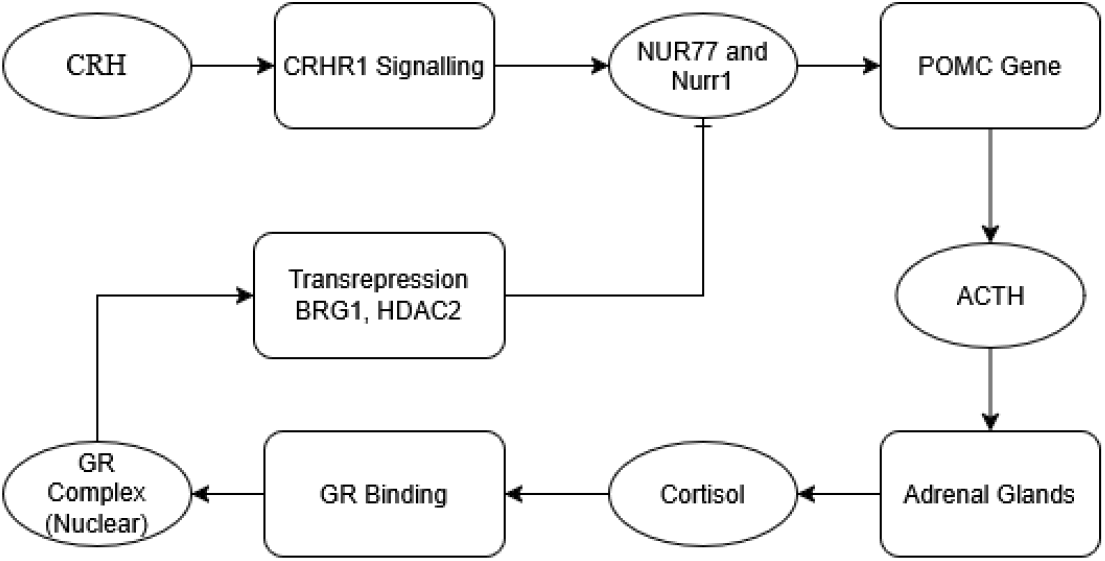
A simplified schematic of how cortisol represses further ACTH production. Cortisol binds to the glucocorticoid receptor (GR), which, following nuclear translocation, can transrepress Nur77/Nurr1-mediated activation of POMC transcription, thereby reducing ACTH synthesis [10, 12].

**Figure 2.**
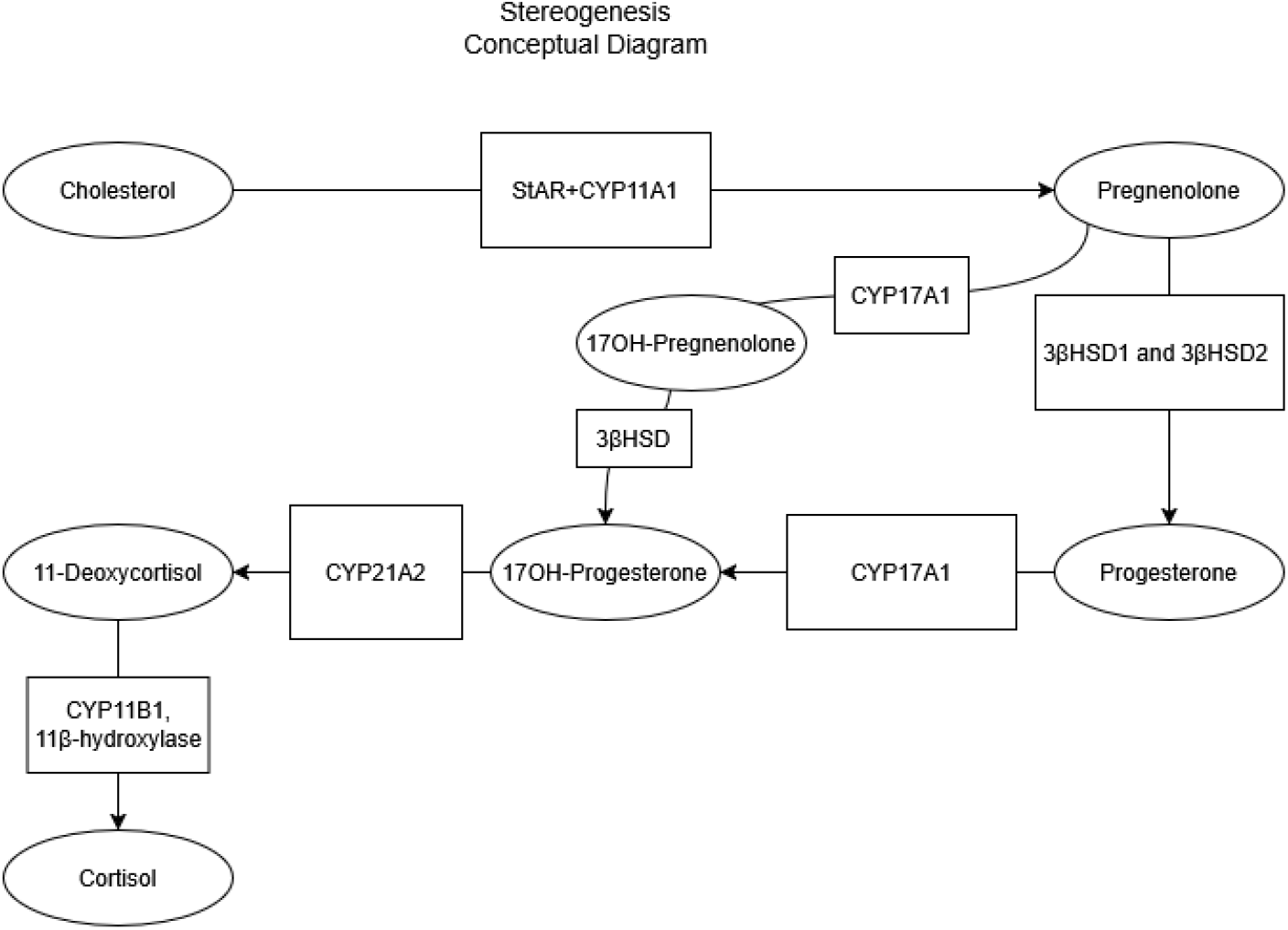
A conceptual graph of the process of steroidogenesis, based on Miller et al [14]. The cascade takes place as a result of ACTH signals communicated via the bloodstream.

**Figure 3.**
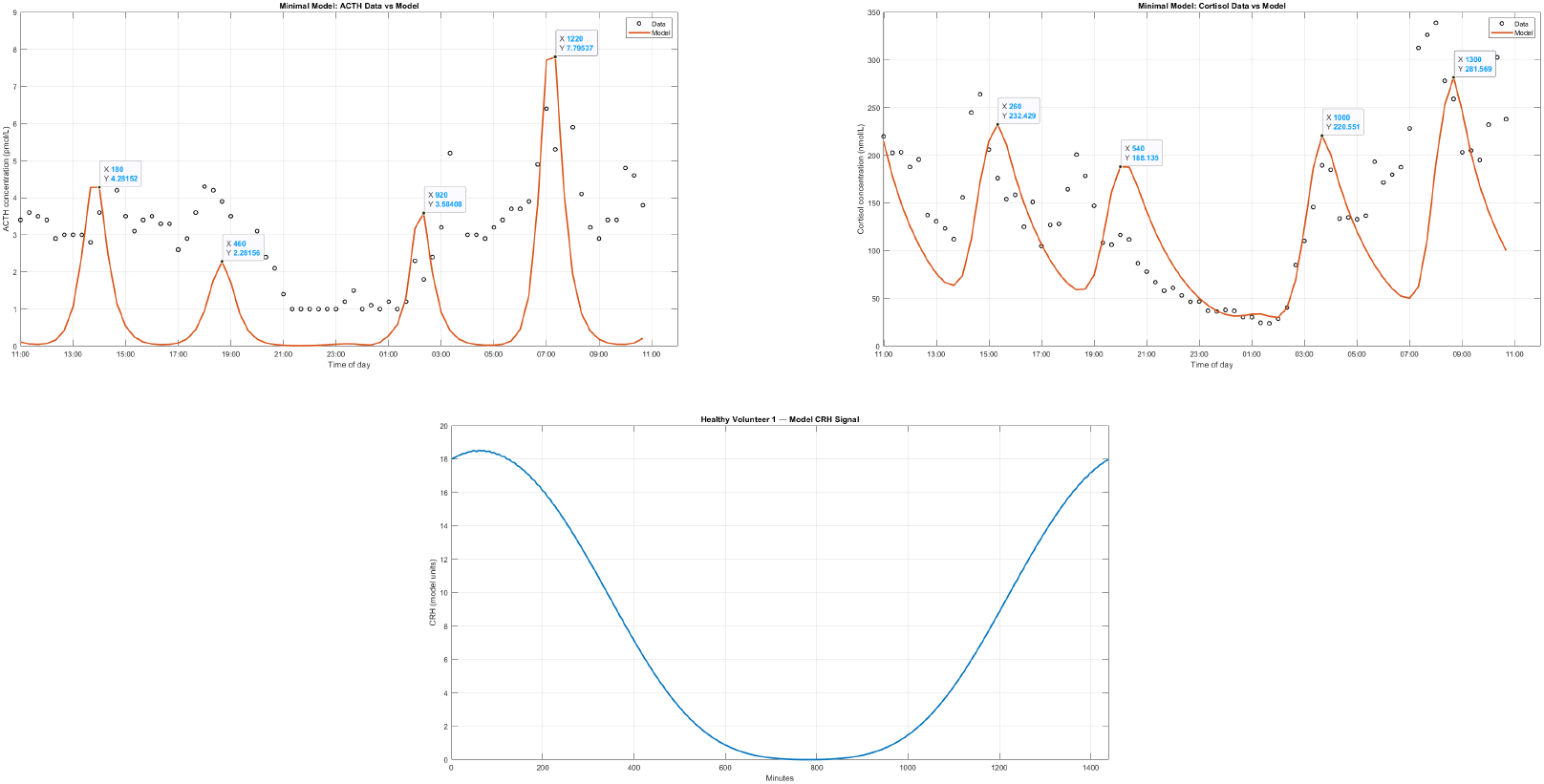
Example fit of Healthy Volunteer 1 using the minimal model. The upper-left panel shows cortisol concentration (nmol/L) over time, with the model trajectory compared with the experimental measurements; the upper-right panel shows the corresponding modelled and measured ACTH concentration (pmol/L). The lower panel shows the circadian CRH input used to drive the model. Time is expressed in minutes. Although the model reproduces the broad structure of the measured profiles, it requires an unrealistically large ACTH-to-cortisol lag to do so.

**Figure 4.**
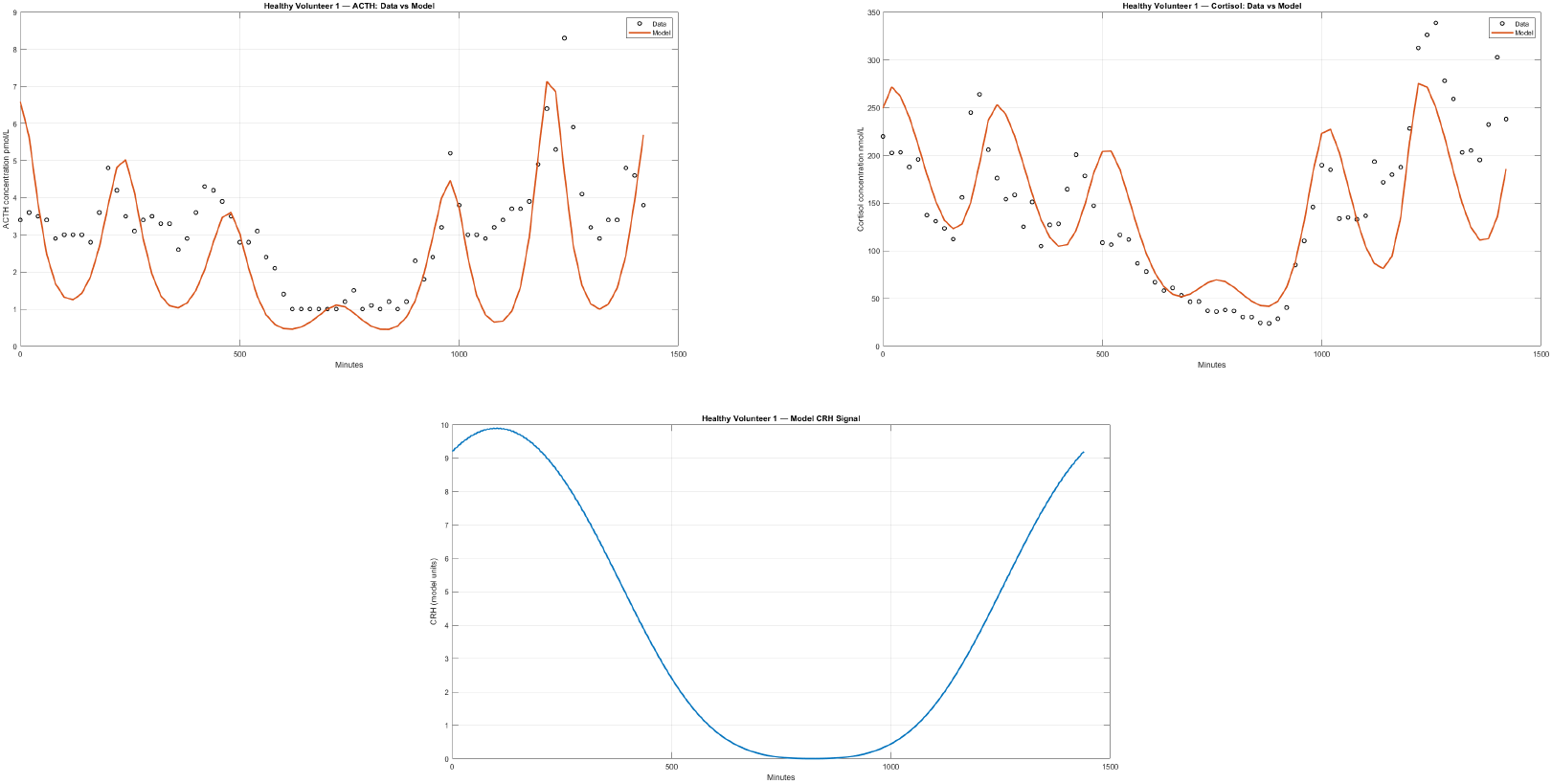
Mechanistic model simulations fitting Healthy volunteer 1 from Upton et al [18]. Fits for cortisol and ACTH are shown as well as the model’s inferred CRH signal.

**Figure 5.**
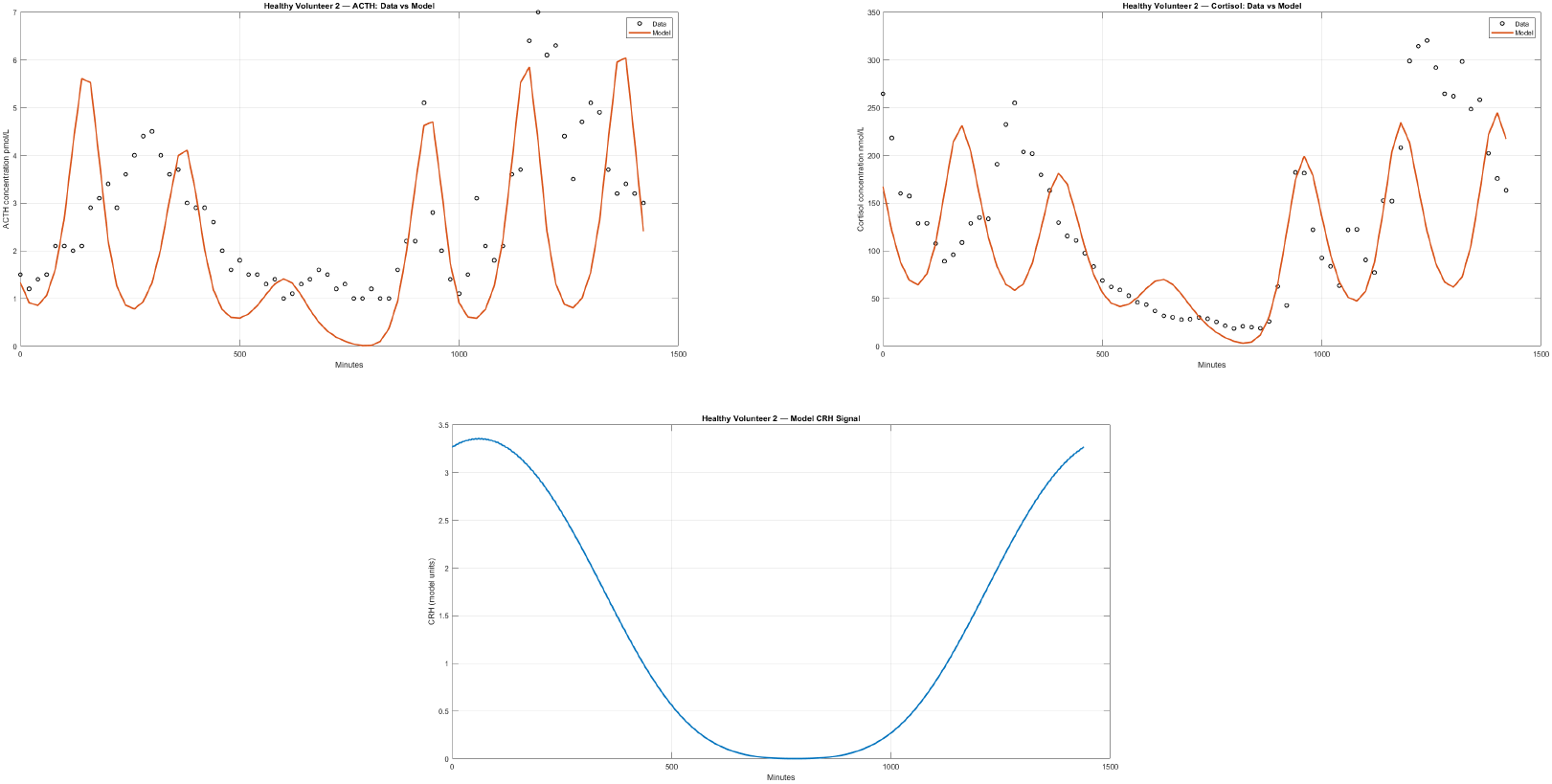
Mechanistic model simulations fitting Healthy volunteer 2 from Upton et al [18]. Fits for cortisol and ACTH are shown as well as the model’s inferred CRH signal

**Figure 6.**
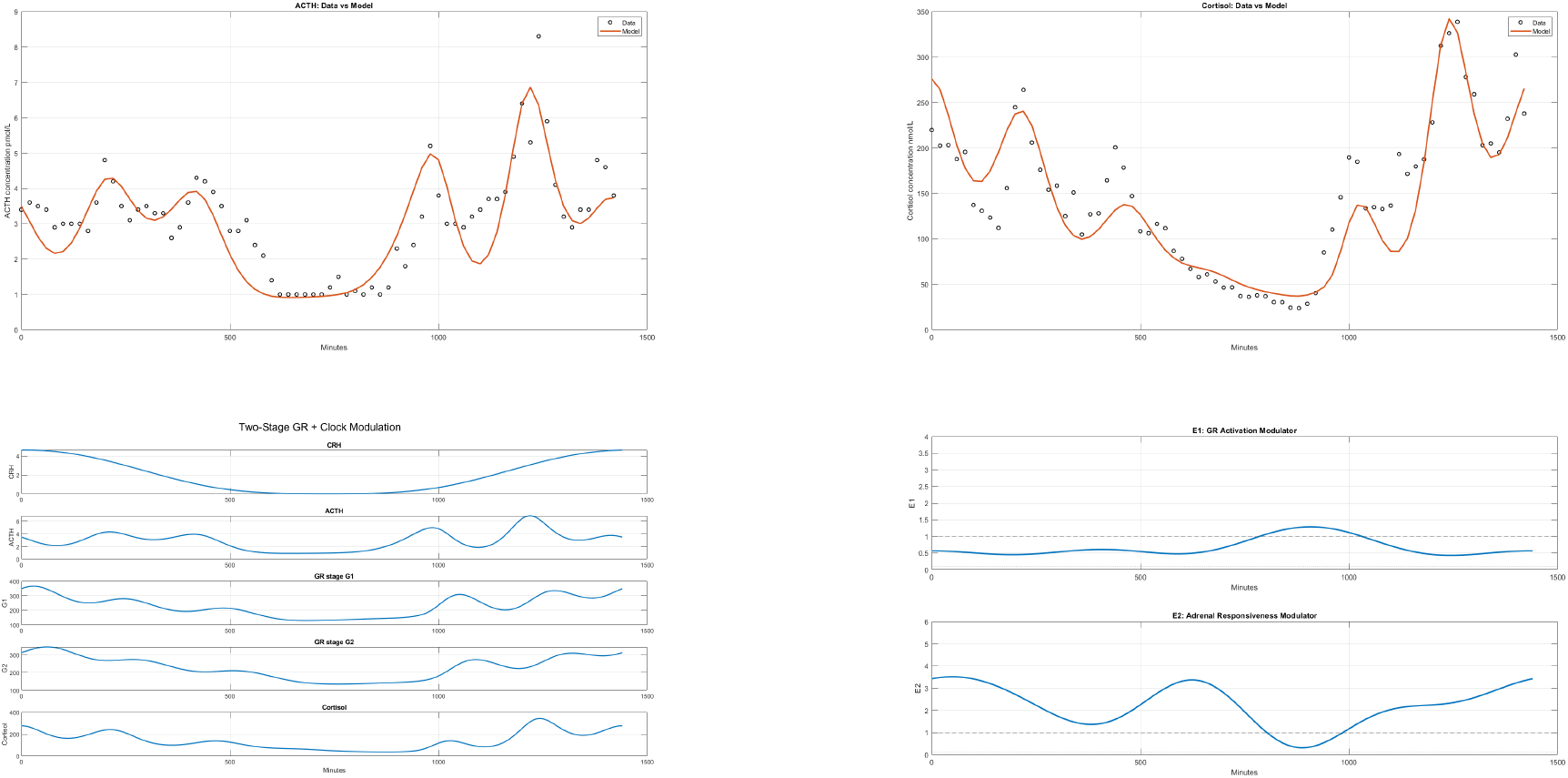
The fit of Volunteer 1 by the modulated model. Here we present all graphs as well as the Fourier modulating series.

**Figure 7.**
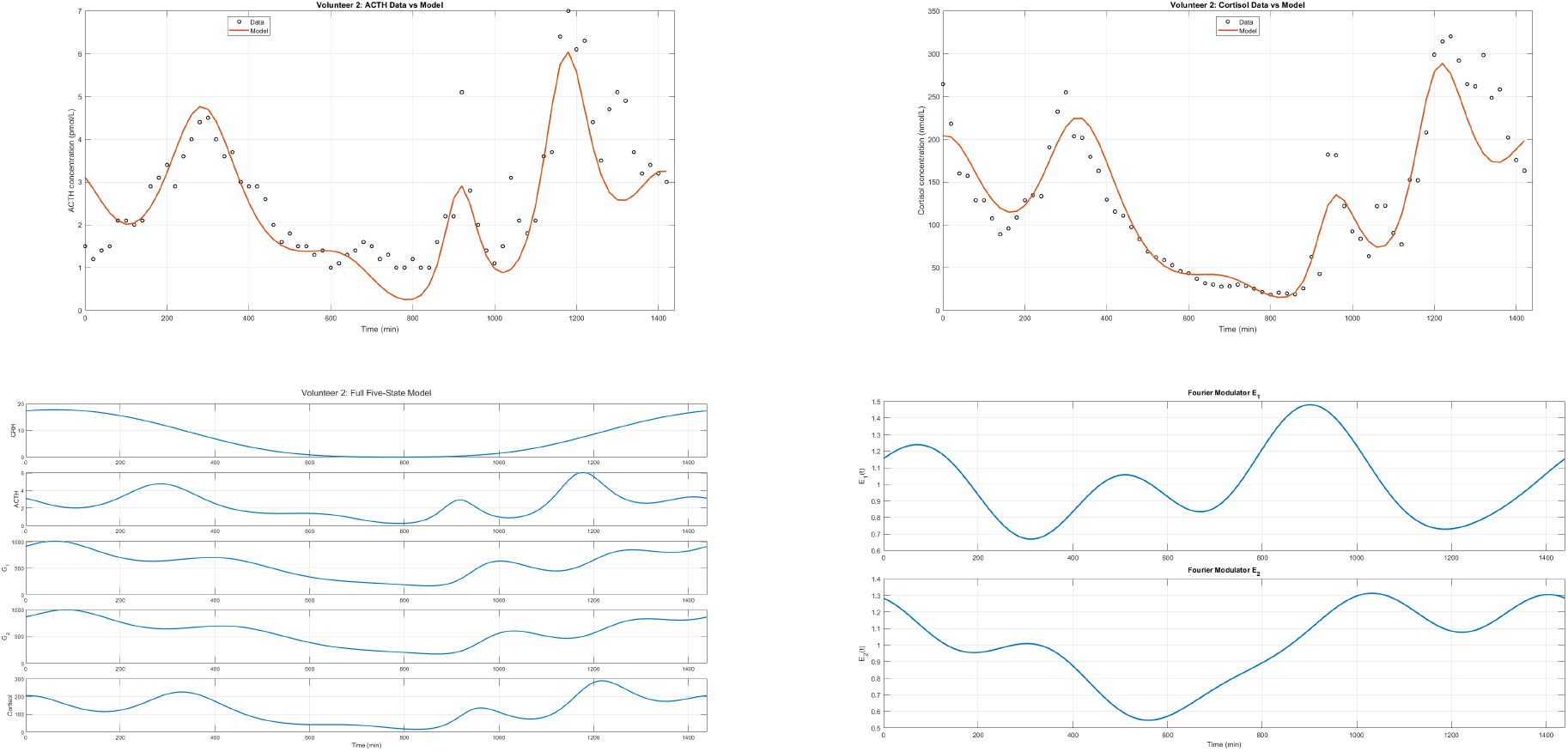
The fit of Volunteer 2 by the modulated model. Here we present all graphs as well as the Fourier modulating series.

**Figure 8.**
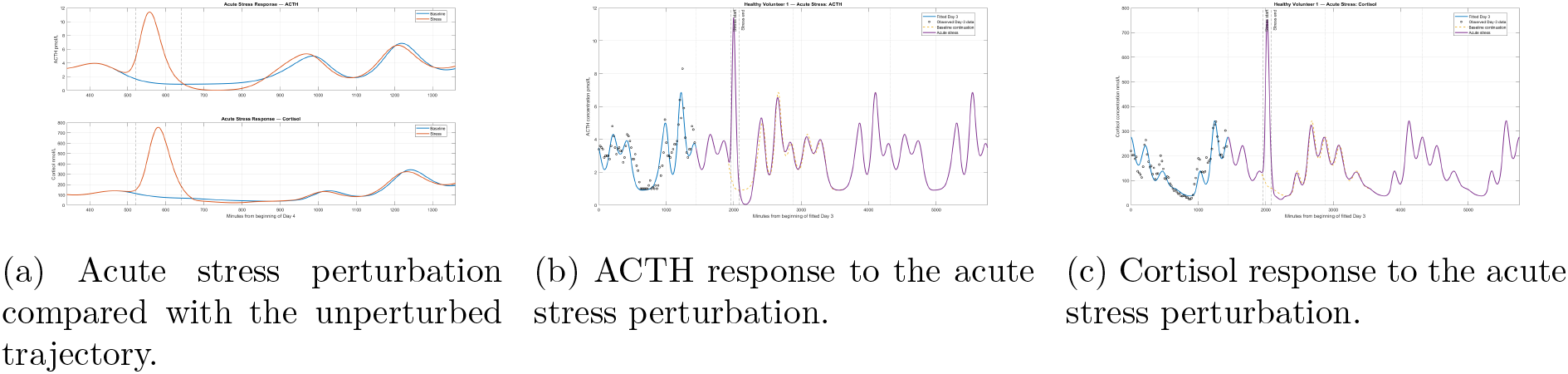
Response of the fitted HPA-axis model to an acute stress perturbation. (a) Acute stress response compared with the corresponding unperturbed trajectory. (b) ACTH dynamics before and after the perturbation. The model is first shown over the interval used for fitting Healthy Volunteer 1 and is then continued forward in time using the fitted parameter values, with and without the acute stress input. (c) Corresponding cortisol dynamics under the same continuation procedure. Following removal of the stress input, the perturbed ACTH and cortisol trajectories progressively return towards the corresponding unstressed trajectories.

Beyond the experimental observation window, the fitted system was integrated forward using the same fitted parameter values. Because *E*_1_(*t*) and *E*_2_(*t*) were constructed from integer harmonics of a 24-h fundamental frequency, the fitted modulatory waveforms repeat intrinsically with a period of 1440 min throughout the extended simulation.

Although this behaviour was observed for most of the parameter sets examined, some simulations displayed substantially different post-perturbation trajectories. An example obtained using the Healthy Volunteer 1 parameterization is shown in Fig. 9.

**Figure 9.**
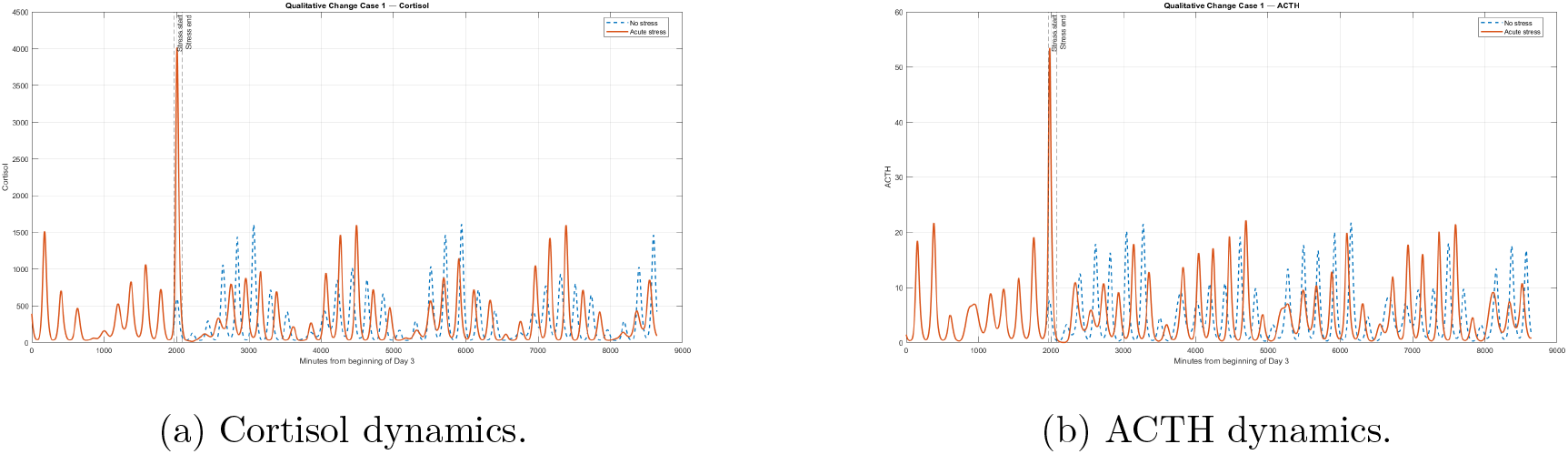
Persistent post-perturbation changes in ACTH and cortisol dynamics following an acute stress pulse for Healthy Volunteer 1.

In this case, the perturbed trajectory did not immediately return to the unstressed trajectory after removal of the stress input. Instead, the subsequent ACTH and cortisol profiles remained visibly separated from the corresponding control trajectories over the simulated interval. Similar qualitative changes were observed for additional parameter sets generated in the neighbourhood of the Healthy Volunteer 1 fit, two representative examples of which are shown in Fig. 10.

**Figure 10.**
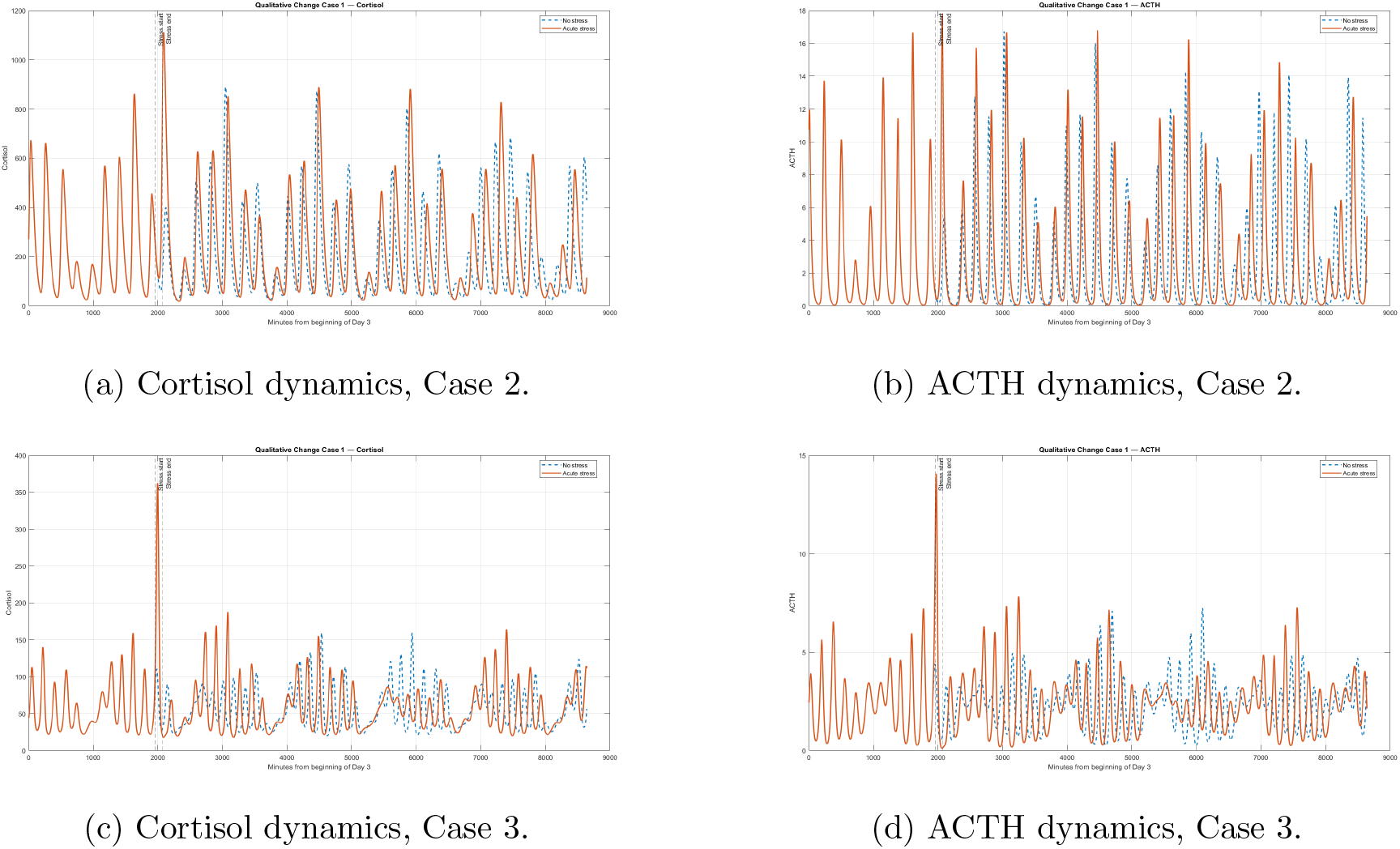
Persistent qualitative changes in the post-stress trajectories for two representative parameter sets generated around the Healthy Volunteer 1 fit.

Additional parameterizations were generated by randomly perturbing the fitted mechanistic parameters of Healthy Volunteer 1 within a restricted neighbourhood of the original fit, with each parameter varied between 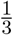 and 2 times its fitted value. These perturbed parameter sets were then simulated forward, and representative cases exhibiting qualitatively different behaviour during the subsequent day were selected for illustration.

These simulations alone do not establish the existence of a bifurcation, because persistent divergence following a finite perturbation may also result from phase resetting or a long transient. They nevertheless indicated that the model could possess parameter regimes in which its qualitative response to stress changes substantially. This motivated a formal bifurcation analysis of the autonomous reduction of the five-state model. Formal bifurcation analysis confirmed that the autonomous model can undergo stress-induced changes in stability for specific combinations of the phenomenological modulatory parameters *E*_1_ and *E*_2_. These parameters represent the constant counterparts of the time-dependent modulatory signals used in the fitted model and were varied relative to their corresponding mean values, *Ē*and *Ē*.

For this analysis, the explicitly time-dependent model was reduced to an autonomous system by replacing *T*_1_(*t*) with its 24-h mean *T*_0_, replacing the periodic modulators *E*_1_(*t*) and *E*_2_(*t*) with constant parameters *E*_1_ and *E*_2_, and representing sustained stress by a constant input *S*; the remaining model structure and fitted parameters were unchanged (see Methods for details).

Continuation of the equilibrium solutions revealed well-defined Hopf bifurcation boundaries in the (*S, E*_1_) and (*S, E*_2_) parameter planes (Fig. 11). Across a substantial region of parameter space, considerably large values of sustained stress could be applied without producing a change in equilibrium stability. However, the critical stress required to reach a Hopf bifurcation depended strongly on the modulatory state of the system.

**Figure 11.**
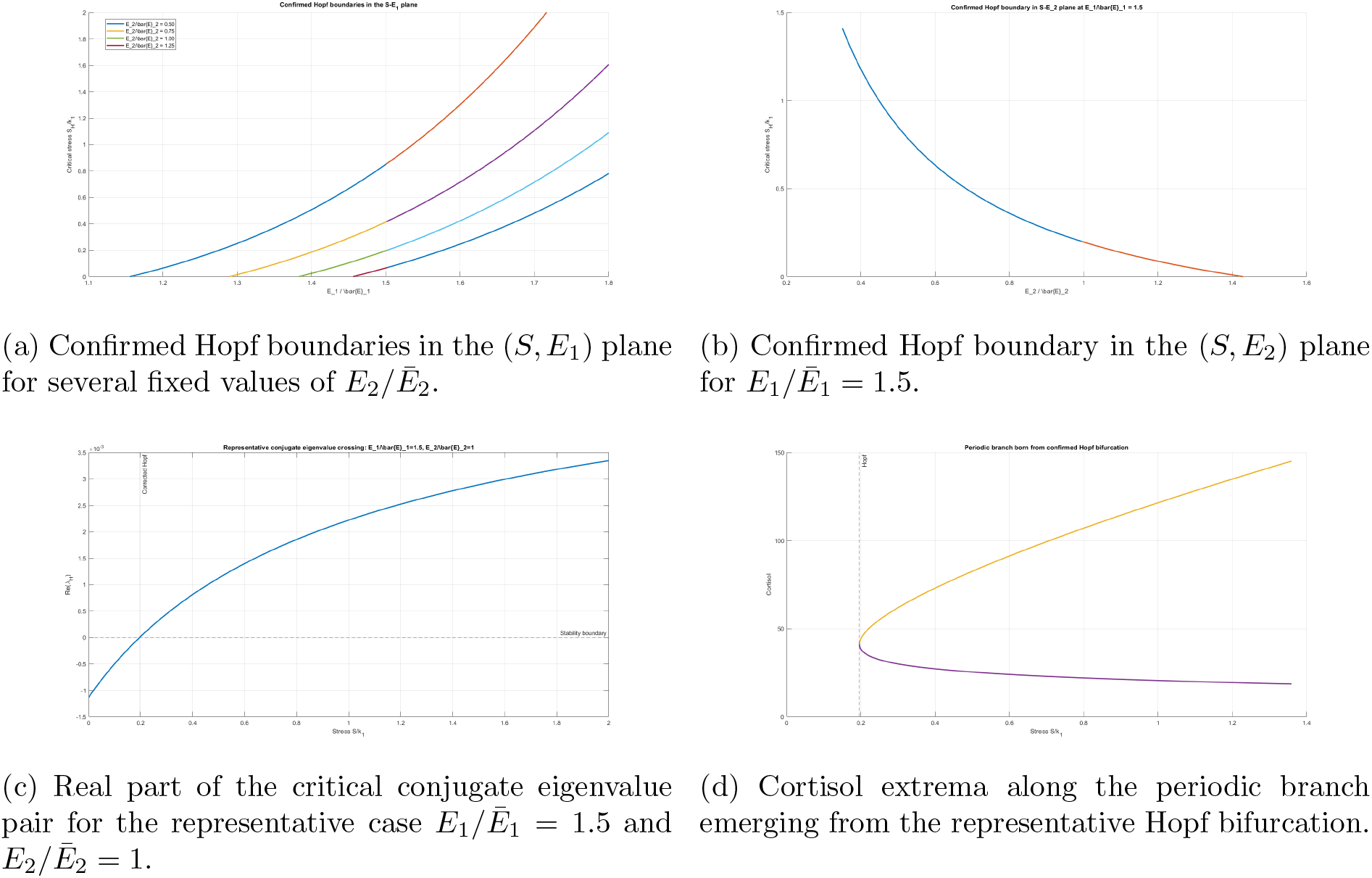
Bifurcation structure of the autonomous HPA-axis model under sustained stress. (a,b) Dependence of the critical stress threshold on the phenomenological modulatory parameters *E*_1_ and *E*_2_. (c) Transversal crossing of the imaginary axis by the critical conjugate characteristic roots at a representative Hopf point. (d) Periodic branch emerging from the corresponding Hopf bifurcation.

The influence of *E*_2_ on the stress threshold was particularly pronounced. For fixed *E*_1_*/Ē*_1_ = 1.5, corrected Hopf points were obtained at

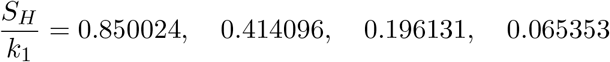

for

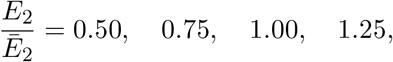

respectively. Thus, over this parameter range, increasing *E*_2_ markedly reduced the magnitude of sustained stress required to destabilize the equilibrium. Conversely, continuation in the (*S, E*_1_) plane showed that increasing *E*_1_ generally shifted the corresponding Hopf boundary towards larger stress values. The resulting curves therefore define sections of a critical surface

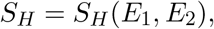

demonstrating that the susceptibility of the autonomous system to sustained stress depends strongly on the balance between the two modulatory factors.

All four corrected Hopf points at *E*_1_*/Ē*_1_ = 1.5 were associated with a transition from zero to two unstable characteristic roots. Thus, in each case, an initially stable equilibrium lost stability as the stress parameter crossed the corresponding critical value. Interestingly, the critical frequency remained almost unchanged across these parameter combinations, with

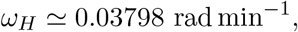

corresponding to

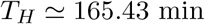

(2.76 h). Variation of *E*_2_ therefore strongly altered the location of the stability boundary while having little effect on the oscillatory timescale selected at the bifurcation.

For example, at *E*_1_*/Ē*_1_ = 1.5 and *E*_2_*/Ē*_2_ = 1.25, the corrected Hopf point occurred at

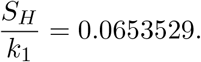

Immediately below this value, the real part of the critical characteristic pair was −8.23 × 10^−6^, whereas immediately above the bifurcation it was 2.14 × 10^−5^. The number of unstable characteristic roots simultaneously changed from zero to two, while the numerically estimated transversality slope was 5.88 × 10^−3^. These results confirm a transversal loss of equilibrium stability through Hopf bifurcation.

The same behaviour is illustrated for the representative case *E*_1_*/Ē*_1_= 1.5 and *E*_2_*/Ē*_2_ = 1 in Fig. 11c. In this case, the corrected bifurcation occurred at

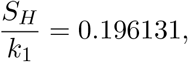

where the real part of the critical conjugate pair crossed zero. Continuation of periodic solutions from this point produced an oscillatory branch whose amplitude increased continuously with sustained stress (Fig. 11d).

The properties and stability of this periodic branch were examined further using its oscillation period and non-trivial Floquet multipliers (Fig. 12). Close to the Hopf point, the period was approximately 165 min and increased gradually with stress, reaching approximately 181 min over the continuation interval examined. The dominant non-trivial Floquet multiplier remained below unity throughout the computed branch and decreased progressively as stress increased, indicating that the periodic solutions remained locally stable and became more strongly attracting away from the bifurcation.

**Figure 12.**
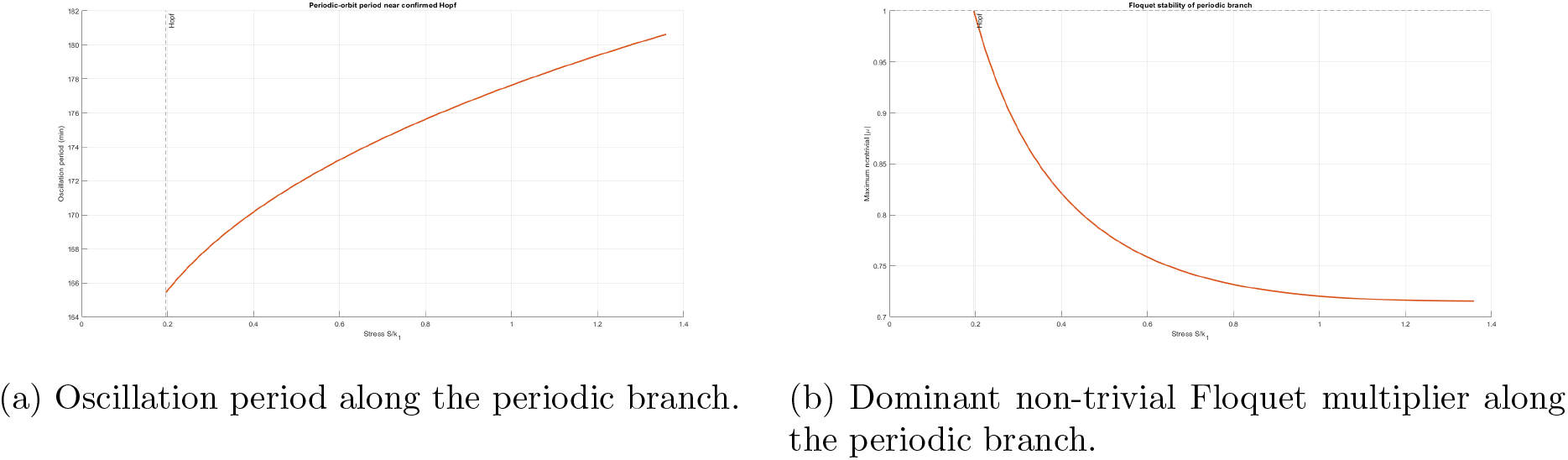
Dynamical properties of the periodic branch emerging from the representative Hopf bifurcation. The oscillation period increases gradually with sustained stress, while the dominant non-trivial Floquet multiplier remains inside the unit circle.

**Figure 13.**
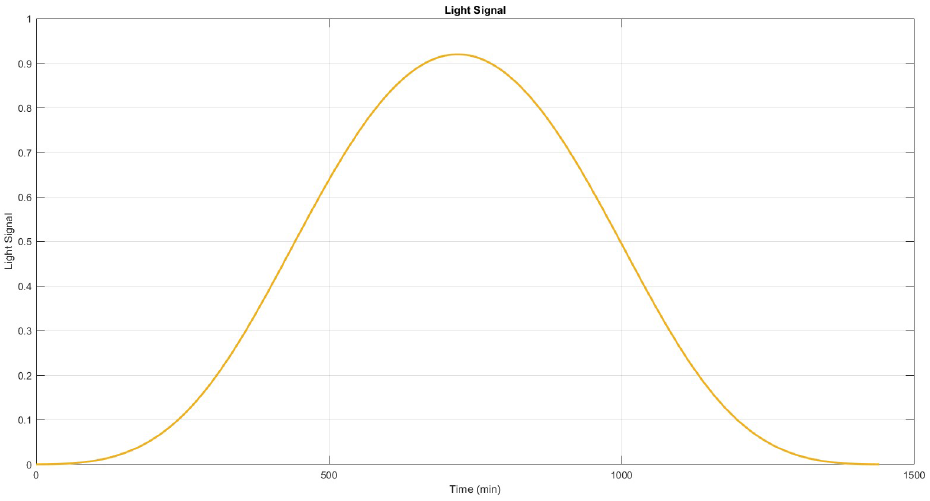
Light Signal over a day starting at 12:00 o’ clock at night and ending at 12:00 o’ clock at night of the following day (1440 minutes).

**Figure 14.**
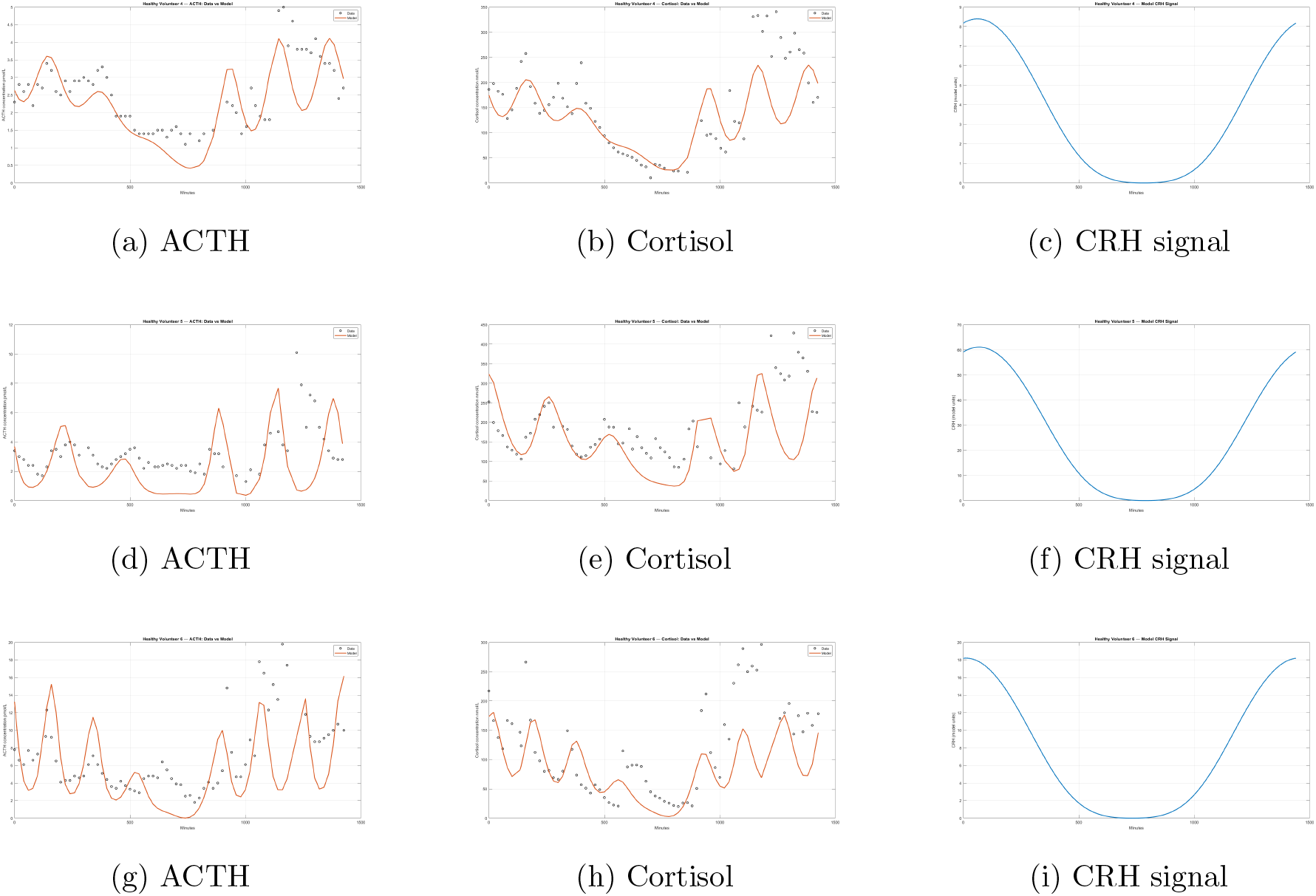
Model fits and CRH signals for Healthy Volunteers 4–6. Rows correspond to Volunteers 4, 5, and 6, respectively. Columns show ACTH data and model predictions, cortisol data and model predictions, and the corresponding model CRH signal.

**Figure 15.**
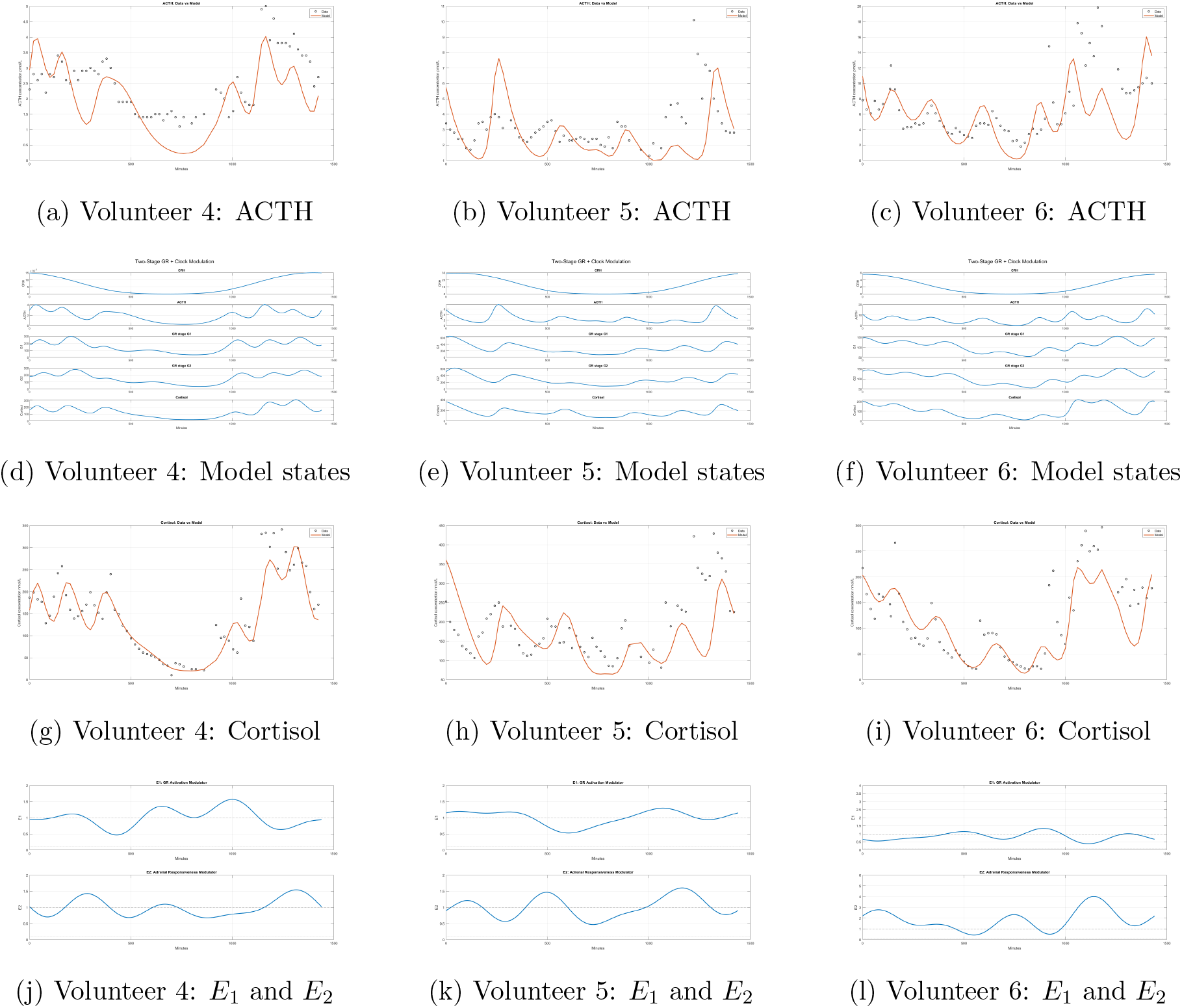
Fourier-modulated model fits and internal model dynamics for Healthy Volunteers 4–6. Columns correspond to Volunteers 4, 5, and 6, respectively. The first row shows ACTH data and model predictions; the second row shows the model state trajectories; the third row shows cortisol data and model predictions; and the fourth row shows the fitted modulatory functions *E*_1_(*t*) and *E*_2_(*t*).

The continuous emergence of a stable, small-amplitude periodic branch from an equilibrium that loses stability through a conjugate-pair crossing, together with the non-trivial Floquet multipliers remaining inside the unit circle, is consistent with a supercritical Hopf bifurcation for the representative parameter set. More generally, the two-parameter continuation results show that the equilibrium remains stable over a broad range of sustained-stress and modulatory conditions, while particular combinations of *E*_1_ and *E*_2_ shift the Hopf boundary substantially and can markedly reduce the stress level required for loss of stability.

## Discussion

The present study investigated the level of dynamical complexity required for a mathematical model of the HPA axis to reproduce simultaneously the temporal characteristics of ACTH and cortisol secretion observed in high-frequency human measurements. Beginning from a minimal three-state formulation, we found that the principal limitation of the reduced architecture was not simply quantitative fitting accuracy. Rather, its feedback structure imposed a mathematical constraint on the phase relationship between ACTH and cortisol. For Hopf-generated ultradian oscillations, the cortisol lag relative to ACTH is constrained to lie between one quarter and one half of the oscillatory period. Consequently, an ultradian period of approximately four hours (as evident from Healthy Volunteer 2 for instance) necessarily produces an ACTH-cortisol lag substantially larger than the approximately 20-min separation observed physiologically. The introduction of two intermediate glucocorticoid-feedback states removes this structural restriction by distributing the phase accumulation required for oscillation among several dynamical stages. The resulting five-state architecture can therefore support ultradian oscillations with both a physiologically appropriate period and ACTH-cortisol phase relationship.

This result provides a dynamical interpretation of why apparently minor additions to an HPA-axis model can produce substantial qualitative changes in its behaviour. The intermediate states *G*_1_ and *G*_2_ should not necessarily be interpreted as two uniquely identifiable molecular species. Rather, they provide an effective representation of intermediate processes separating circulating cortisol from its eventual inhibitory effect on pituitary ACTH secretion. Their mathematical role is nevertheless important: each intermediate dynamical stage contributes additional phase delay to the negative-feedback loop without requiring this phase to appear directly as an excessively large ACTH-cortisol lag. Thus, the improvement obtained with the five-state system results not merely from adding additional fitting parameters, but from changing the phase structure of the feedback architecture.

The unmodulated five-state model reproduced the principal features of the measured ACTH and cortisol profiles more successfully than the minimal formulation and produced physiologically appropriate ACTH-cortisol timing in most of the fitted volunteers. The addition of the time-dependent modulatory functions *E*_1_(*t*) and *E*_2_(*t*) produced a further substantial improvement in the agreement between model and data. These functions were introduced as phenomenological representations of time-varying processes that are omitted from the core model. *E*_1_(*t*) may therefore be interpreted as an effective modulation of glucocorticoid feedback, whereas *E*_2_(*t*) represents effective variation in adrenal responsiveness to ACTH. Their biological interpretation is intentionally non-unique. Processes including receptor regulation, clock-dependent signalling, cAMP and StAR dynamics, unmeasured central input and other physiological influences may all contribute to the modulation inferred from the hormone profiles.

This phenomenological interpretation is particularly important because the modulatory signals are inferred indirectly from ACTH and cortisol measurements rather than measured independently. Their fitted waveforms should therefore not be regarded as direct reconstructions of specific molecular signals. Instead, they identify the magnitude and temporal structure of additional modulation required by the measured hormone profiles once the principal HPA feedback architecture has been specified. The smooth and constrained nature of the inferred functions also provides a potential target for future models in which the Fourier representation is replaced by an explicit biological oscillator or regulatory subsystem.

Stress-response simulations showed that the model is capable of homeostatic recovery following acute perturbation for the fitted Healthy Volunteer 1 parameterization. In most simulations the perturbed trajectory returned towards the corresponding unstressed dynamics after the stress input was removed. In a smaller subset of parameter combinations, however, transient stress produced persistent differences between the perturbed and control trajectories over the simulated interval. Such simulations alone cannot distinguish a genuine change of attractor from phase resetting or a prolonged transient. Their importance in the present study was therefore primarily diagnostic: they indicated parameter regions in which the qualitative response of the model to stress changed and motivated a formal bifurcation analysis.

The autonomous reduction of the five-state system confirmed that sustained stress can induce a Hopf bifurcation, with the critical stress threshold depending strongly on the effective modulatory state of the system. In particular, increasing *E*_1_ generally shifted the Hopf boundary towards larger values of sustained stress, whereas increasing *E*_2_ markedly reduced the stress required for destabilization. This behaviour has a natural interpretation within the structure of the model. Stronger effective glucocorticoid feedback tends to oppose the destabilizing effect of sustained activation, whereas increased adrenal responsiveness increases the effective gain between ACTH and cortisol and can move the feedback system closer to an oscillatory instability. Importantly, the bifurcation threshold was considerably more sensitive to these modulatory conditions than the oscillatory frequency itself: across the parameter combinations examined, the critical period remained close to 165 min even though the stress threshold changed substantially. Continuation of the periodic solutions arising from a representative Hopf point further showed that the oscillatory branch remained locally stable over the computed interval, while its period increased gradually with sustained stress. The continuous emergence of a stable, small-amplitude periodic branch from the equilibrium is consistent with a supercritical Hopf bifurcation for this parameter set. These results demonstrate that the model can undergo a qualitative transition from a stable equilibrium to a stable oscillatory regime through variation of sustained stress and effective systemic modulation. They do not, however, by themselves establish bistability or hysteresis, which would require demonstration of coexisting attractors and dependence of the asymptotic state on the direction or history of parameter variation.

The bifurcation results also suggest a way of interpreting inter-individual or time-dependent differences in HPA-axis susceptibility without assuming that stress intensity alone determines the dynamical response. Within the model, the same magnitude of sustained stress can lie either below or above the stability boundary depending on the prevailing values of the modulatory factors. The relevant control variable is therefore not stress in isolation but its interaction with the regulatory state of the HPA axis. This provides a mathematical mechanism through which changes in feedback sensitivity or adrenal responsiveness could alter the response to otherwise comparable external perturbations. At present, however, the model does not establish a direct correspondence between particular regions of the bifurcation diagram and specific clinical disorders. Such an interpretation would require parameterization and testing against data obtained from relevant patient populations.

Several limitations should therefore be emphasised. First, the model was calibrated using hormone profiles from five healthy volunteers, and the same profiles were used to assess the quality of the fits. Second, CRH and the intermediate feedback states were not measured directly and are consequently latent model variables. Third, the Fourier modulators introduce additional flexibility and are not uniquely identifiable with specific biological mechanisms. Fourth, the formal Hopf analysis was performed on an autonomous reduction in which the circadian CRH input and periodic modulators were replaced by constant quantities. The resulting bifurcation structure therefore characterises the underlying autonomous feedback architecture rather than proving an equivalent bifurcation of the complete periodically forced model. Finally, the imposed stress term is phenomenological and does not attempt to reproduce the detailed neural pathways through which different classes of physical or psychological stressors activate the HPA axis.

Future work should address these limitations by examining the stability of the complete periodically forced system, replacing the phenomenological modulators with explicit regulatory oscillators where suitable data permit, and testing the resulting model against additional high-frequency endocrine profiles. Measurements capable of constraining latent components of the feedback pathway would be particularly valuable for distinguishing between alternative mechanistic interpretations of the inferred modulation. Application of the model to pathological endocrine profiles could subsequently determine whether the bifurcation structure identified here corresponds to clinically relevant transitions in HPA axis dynamics.

In summary, the present results show that reproducing human HPA axis dynamics requires attention not only to hormone concentrations but also to the phase structure of the underlying feedback loop. The minimal three-state architecture imposes a phase constraint that is incompatible with the simultaneous occurrence of a physiologically realistic ACTH-cortisol lag and ultradian period, whereas the five-state architecture removes this restriction through intermediate feedback dynamics. When supplemented by constrained time-dependent modulation, the model reproduces a range of healthy human hormone profiles and exhibits stress-dependent Hopf bifurcations whose location depends strongly on the regulatory state of the system. Together, these results provide a dynamical framework for investigating how physiological modulation and sustained stress may interact to produce qualitative changes in HPA-axis behaviour.

## Methods

### Minimal model

We begin with a deliberately minimal model consisting of three state variables: CRH, ACTH, and cortisol. The corresponding dynamics are described by the following system of delay differential equations:

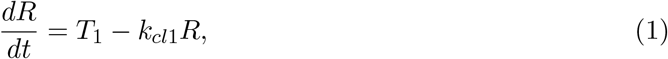

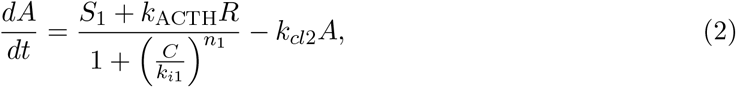

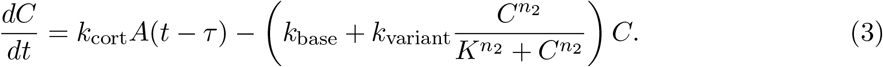

The model variables and parameters, together with their definitions and units, are summarized in Table 1. The variable *R* denotes the CRH signal, light induced, *A*, ACTH and *C* cortisol.

**Table 1:**
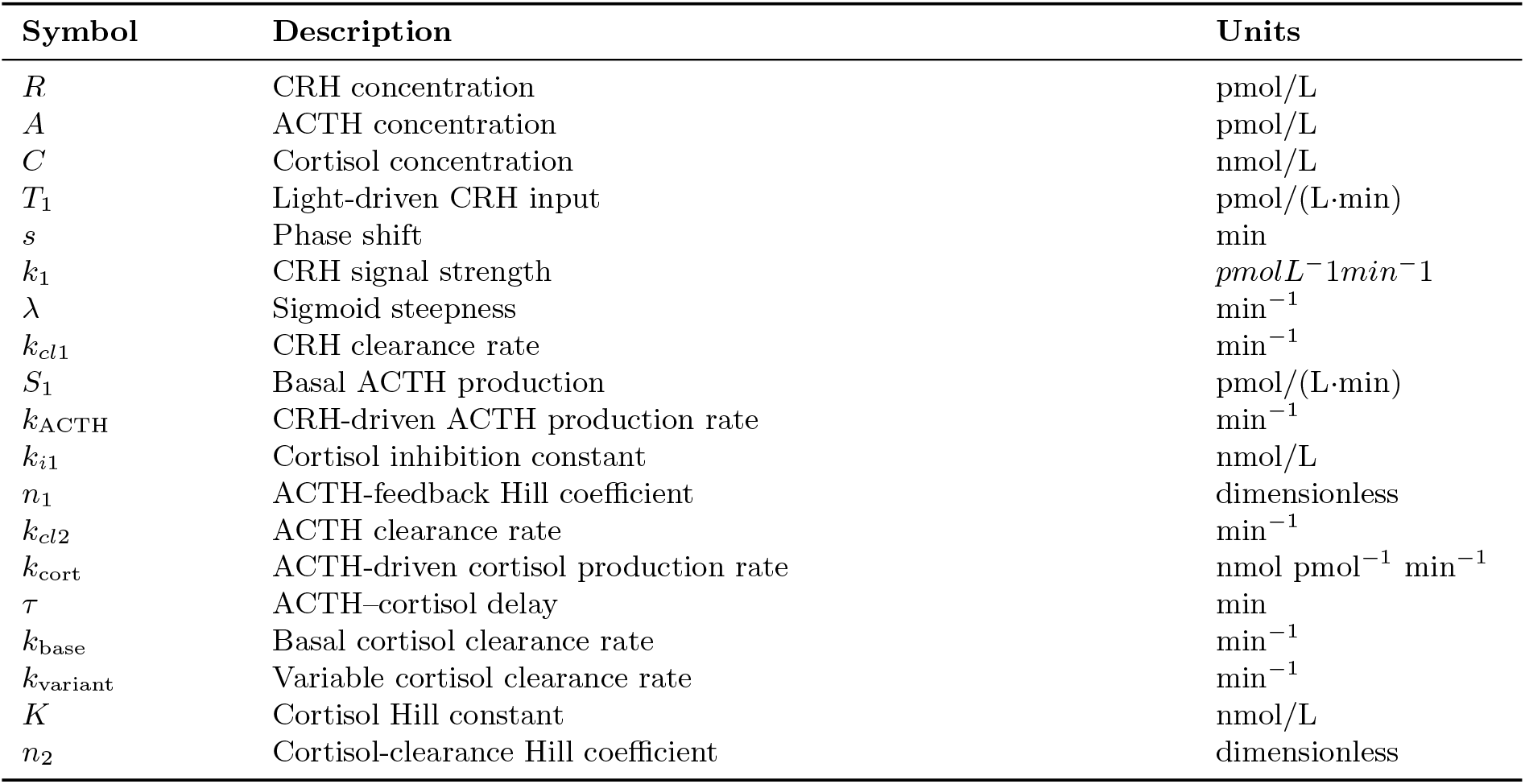
Variables and parameters of the minimal HPA-axis model.

| Symbol | Description | Units |
| --- | --- | --- |
| $R$ | CRH concentration | pmol/L |
| $A$ | ACTH concentration | pmol/L |
| $C$ | Cortisol concentration | nmol/L |
| $T_1$ | Light-driven CRH input | pmol/(L·min) |
| $s$ | Phase shift | min |
| $k_1$ | CRH signal strength | $\text{pmol L}^{-1} \text{min}^{-1}$ |
| $\lambda$ | Sigmoid steepness | $\text{min}^{-1}$ |
| $k_{cl1}$ | CRH clearance rate | $\text{min}^{-1}$ |
| $S_1$ | Basal ACTH production | pmol/(L·min) |
| $k_{\text{ACTH}}$ | CRH-driven ACTH production rate | $\text{min}^{-1}$ |
| $k_{i1}$ | Cortisol inhibition constant | nmol/L |
| $n_1$ | ACTH-feedback Hill coefficient | dimensionless |
| $k_{cl2}$ | ACTH clearance rate | $\text{min}^{-1}$ |
| $k_{\text{cort}}$ | ACTH-driven cortisol production rate | nmol pmol$^{-1}$ min$^{-1}$ |
| $\tau$ | ACTH–cortisol delay | min |
| $k_{\text{base}}$ | Basal cortisol clearance rate | $\text{min}^{-1}$ |
| $k_{\text{variant}}$ | Variable cortisol clearance rate | $\text{min}^{-1}$ |
| $K$ | Cortisol Hill constant | nmol/L |
| $n_2$ | Cortisol-clearance Hill coefficient | dimensionless |

**Table 2:**
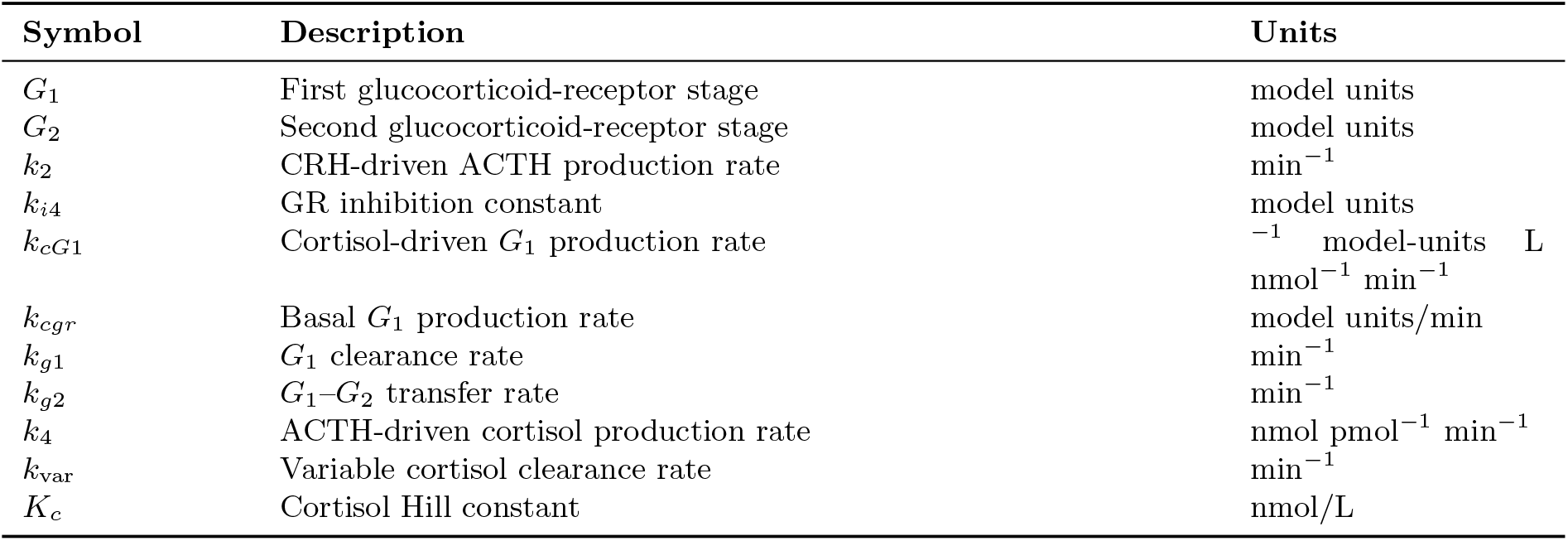
Additional variables and parameters introduced in the five-dimensional mechanistic HPA-axis model.

For the determination of the light signal, several candidate functions were explored, and we selected the present form because it produced a smoother daily waveform than simpler alternatives. The combination of sigmoid and squared sinusoidal terms was also mathematically convenient, since these functions are continuous, continuously differentiable, and smooth, while readily accommodating the time-shift parameter determined by the measurement onset time of each dataset.

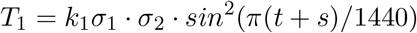

where,

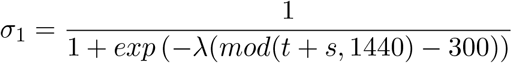

and,

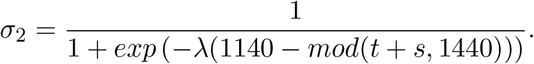

The parameter *s* simply denotes the phase shift, because measurement data begins during different times of the day while *k*_1_ determines the strength of the signal.

The CRH signal represents an idealized light/dark day driver where the signal is near zero during the dark hours of the day and increases in response to light. For rodents and other nocturnal animals the *CRH* signal needs to be inverted. Parameter *s* represents a phase shift that might appear in data since the model’s *zero* is taken to be 12 o clock, midnight. Graphically then the chosen light signal is shown below.

In the case of cortisol we introduce a Hill function to properly capture the fact that its clear-ance is not constant but rather varies depending on cortisol levels. This can be demonstrated by our previous work [16] as well as from Lacerda et al [17]. We achieve the varying rate by using a Hill function.

Cortisol clearance is represented by the sum of a basal component, *k*_base_, and a concentration-dependent component, *k*_variant_. The latter is modulated by a Hill function, so that the effective clearance rate is low when *C* ≪ *K*, increases around the threshold *C* ≈ *K*, and approaches a maximum value as *C* becomes large. The exponent *n*_2_ controls the steepness of this transition. The fitting code was free to choose the exponent, so it is different in each simulation. A similar Hill-type formulation is used to represent the inhibitory effect of cortisol on ACTH production. As cortisol increases, the inhibitory term becomes stronger, thereby reducing ACTH synthesis. The Hill exponent *n*_1_ determines the steepness of this feedback response and is estimated during parameter fitting.

Finally, we provide a table of parameters used in the minimal model. Subsequent tables will only include parameters that do not appear here, as most parameters are denoted in exactly the same way from model to model.

Parameters *λ* and *s* did not participate in the fitting process, as *λ* determines the steepness of the light signal curve and was set to 0.0075 for all simulations, while *s* is the phase shift and is determined by the measurement data, i.e. when each volunteer’s measurements begun during the day.

### The full mechanistic model

To solve the lag problem we introduce a 5 dimensional mechanistic model. As shown in the supplementary materials, Appendix B, this model can indeed produce the correct lag between ACTH and cortisol. Essentially we add to the previous minimal model two extra stages which are related to how the glucocorticoid receptors act as intermediaries between cortisol and the pituitary. These two extra stages provide us with the proper machinery to allow the model to produce the oscillations with the right 10–20 minute lag. Consider the following system of delay differential equations.

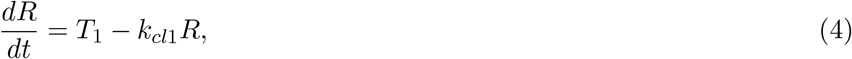

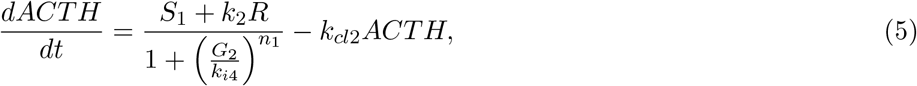

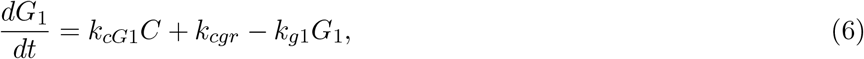

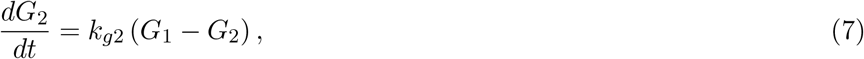

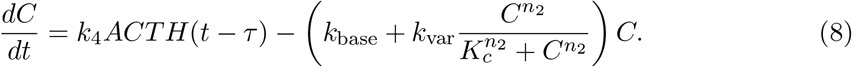

We have added the two extra state variables of *G*_1_ and *G*_2_. Unlike previous models [6],[7] we just use the state parameter *G*_2_ within a Hill function with exponent *n*_1_ to determine ACTH’s response to cortisol. During our research we tried to use the dimer *G*_2_ × *C* as well as other possible architectures but this is the one that is in best agreement with the data. Taking into account the fact that receptors replenish themselves, we also include a basal production rate *k*_*cgr*_ for the *G*_1_ state equation. This is a similar logic with the one appearing in both Gupta et al[6] and Walker et al[7] GR equations, though rather than using a Hill function gated state function, we break GR saturation mechanics into two stages. If one analyzes the mechanistic model as we did for the minimal model in Appendix A, it becomes evident that the addition of two intermediate feedback states is sufficient to remove the phase constraint identified in the minimal architecture. This is important because the phase relationship between ACTH and cortisol is a physiological constraint on the model. An architecture that cannot reproduce the observed lag cannot provide an adequate dynamical representation of the HPA axis.

Here we provide a table for all variables and parameters the extended 5-dimensional model introduces. Any parameter not explicitly written here can be found in table 1

### Mechanistic model with added modulation

Finally, we refine the mechanistic model by introducing two time-dependent modulatory functions, *E*_1_(*t*) and *E*_2_(*t*). These functions are not intended to represent specific biological pathways individually. They are intended to provide phenomenological representations of the net time-varying regulation omitted from the core model. Such regulation may arise from processes including clock-gene-dependent effects, variation in glucocorticoid-feedback transduction, cAMP/StAR availability, and other unmeasured rhythmic influences, as discussed in the introduction. The Fourier modulators therefore represent one possible effective waveform that an underlying biological regulatory system would need to provide.

Each modulator is constructed from a four-harmonic Fourier waveform with a fundamental period of 24 h. We first define the corresponding raw waveform as

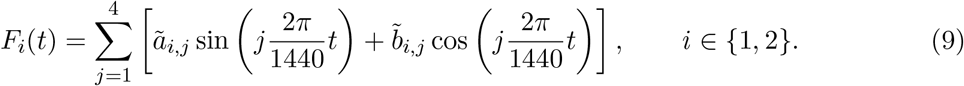

Thus, the harmonic frequencies are fixed at the first four integer multiples of the 24-h fundamental frequency and are not independently fitted.

MATLAB’s particle-swarm algorithm generates candidate values for the harmonic coefficients together with parameters controlling the admissible range of each modulator. For each candidate, the harmonic coefficients first define a raw four-harmonic waveform *F*_*i*_(*t*). The minimum and maximum of this waveform over one 24-h cycle are then calculated, and the waveform is affinely transformed into the fitted interval [*E*_*i*,min_, *E*_*i*,max_]. This transformation introduces the constant offset required to keep the resulting modulator positive, while a separate fitted span parameter *q*_*i*_ controls the amplitude of modulation through

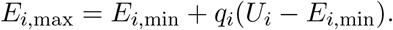

The transformed waveform, rather than the raw Fourier series, is then used in the differential equations, where *U*_1_ = 4 and *U*_2_ = 6. Consequently,

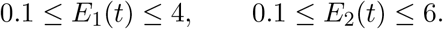

If *F*_*i*,min_ and *F*_*i*,max_ denote the minimum and maximum values of the raw waveform over one 24-h cycle, the modulator used in the differential equations is

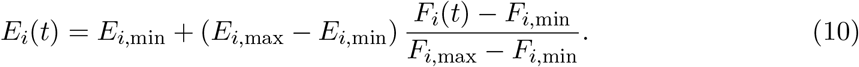

This affine transformation preserves the Fourier structure of the waveform. Consequently, each *E*_*i*_(*t*) remains mathematically equivalent to a constant term plus four sine and four cosine components, while every candidate waveform generated during optimization remains within its prescribed admissible range.

Both *E*_1_(*t*) and *E*_2_(*t*) are dimensionless, since they multiplicatively modulate parameters whose physical dimensions are already defined. The complete model therefore becomes

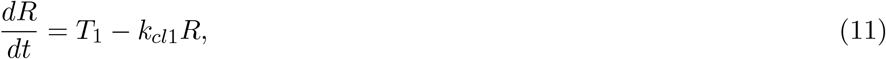

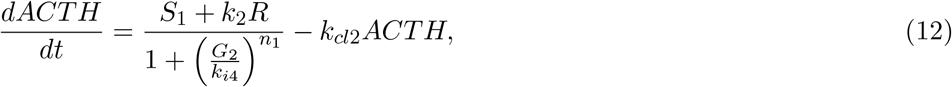

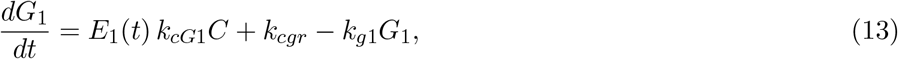

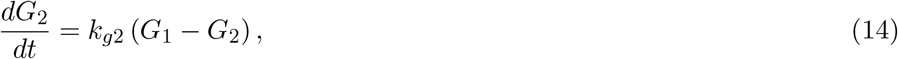

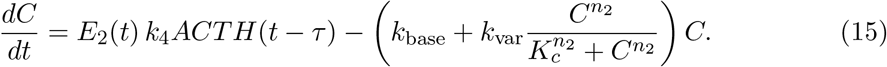

In this formulation, *E*_1_(*t*) represents effective time-dependent modulation of glucocorticoid-feedback transduction, whereas *E*_2_(*t*) represents effective modulation of adrenal responsiveness to ACTH, potentially incorporating processes such as cAMP and StAR regulation. These interpretations are deliberately phenomenological, since the fitted functions may reflect the combined influence of several unmeasured regulatory processes.

The flexibility of the modulatory functions was deliberately restricted during parameter estimation. Their purpose was not to provide arbitrary time-dependent forcing capable of tracking the measured hormone profiles directly. For that reason we used the minimum values, capable of improving the objective *J* discussed in the parameter estimation section as related to the simple mechanistic model. Initial tests used the range [0.1, 2] for both modulators. Where this did not improve the objective for some volunteers, the admissible upper bounds were increased to 4 for *E*_1_ and 6 for *E*_2_, which were the smallest bounds found to provide improvement.

Each modulator introduces ten optimization variables: eight raw Fourier shape coefficients, *ã*_*i*,1_, …, *ã*_*i*,4_ and 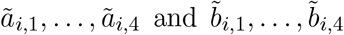, together with the lower-bound parameter *E*_*i*,min_ and the span parameter *q*_*i*_. The two modulators therefore introduce 20 optimization variables in addition to the 17 parameters of the mechanistic model, giving a total optimization vector of 37 variables.

### Parameter estimation

Model parameters were estimated using MATLAB’s particle-swarm optimization algorithm. Rather than minimizing a simple mean-squared error, the objective function was designed to capture several characteristics of the observed ACTH and cortisol profiles, including the timing and number of physiological peaks, the timing of the principal trough, the overall amplitude range, the mean hormone concentration, pointwise agreement with the measurements, and the smoothness of the resulting trajectory.

Each candidate parameterization was simulated for three consecutive days using MATLAB’s `dde23` delay-differential-equation solver. Only the final simulated day was compared with the experimental data, allowing transients associated with the constant DDE history function to decay before evaluation of the fit.

The ACTH objective was defined as

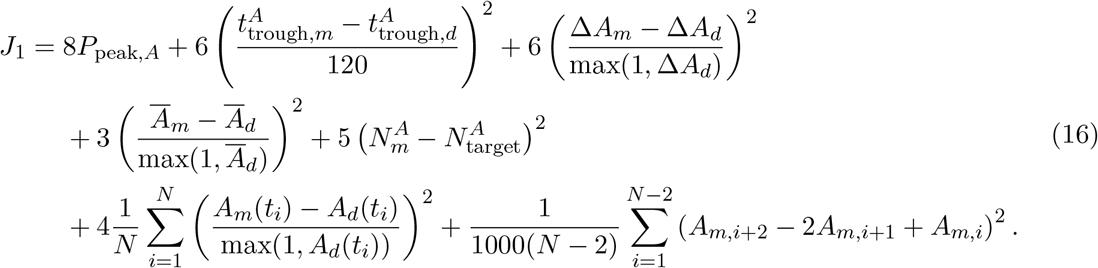

Similarly, the cortisol objective was

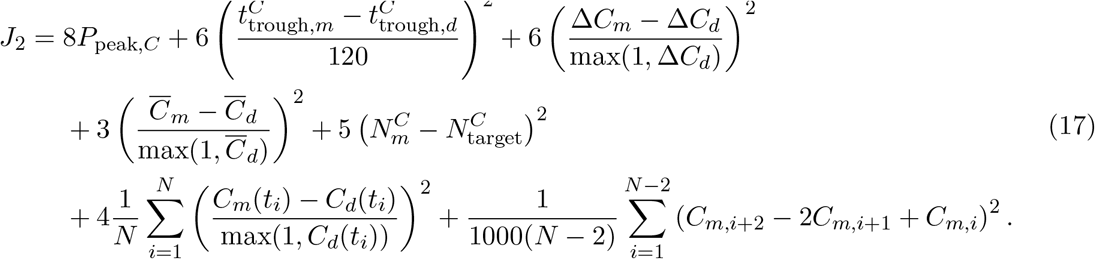

Here, the subscripts *m* and *d* denote model and experimental data, respectively. The quantitiesΔ*A* andΔ*C* denote the corresponding profile ranges, while 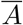 and 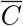 denote their mean values. The quantities 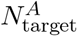and 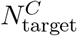 are the numbers of physiological ACTH and cortisol peaks specified from inspection of the corresponding experimental profiles prior to optimization, whereas 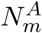 and 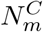 are the numbers of peaks generated by the model.

The terms *P*_peak,*A*_ and *P*_peak,*C*_ quantify the timing mismatch between the specified physiological peak locations and the corresponding model-generated peaks. Peaks were matched chronologically, with the timing penalty calculated from the normalized squared differences between matched peak times. The separate peak-number term penalizes missing or additional model-generated peaks.

The total fitting objective was

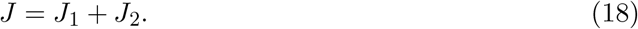

For the full model, optimization was performed by alternating between the 17 mechanistic parameters and the 20 modulation-related variables defining *E*_1_(*t*) and *E*_2_(*t*), rather than optimizing all 37 variables simultaneously.

### Stress-response and bifurcation analysis

To examine the response of the model to external perturbations, we introduced stress directly into the CRH equation. A physiologically plausible regulatory model would be expected to exhibit homeostatic recovery following transient perturbations over a substantial region of parameter space. We therefore performed a series of stress-response experiments both to assess this homeostatic behaviour and to identify parameter regimes in which qualitative changes in the dynamics might occur.

For acute stress, we introduced the term

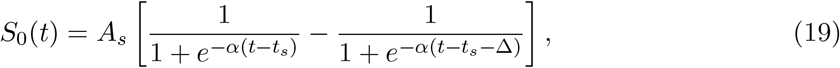

where *A*_*s*_ denotes the stress amplitude, *t*_*s*_ the onset time,Δ the duration of the perturbation, and *α* controls the sharpness of its onset and termination. Stress pulses with durations ranging from several minutes to approximately two hours were examined. Sustained-stress experiments were performed using the same functional form, with the onset and termination times chosen so that the stress input remained active over approximately one day.

We additionally considered repeated stress exposure by introducing a sequence of stress

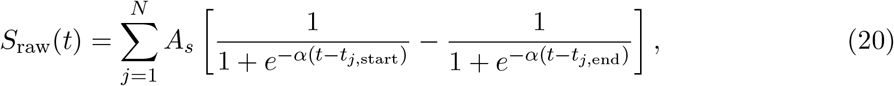

with pulse onset and termination times distributed throughout the simulated day. These experiments were performed both using the fitted parameter set of Healthy Volunteer 1 and using additional parameter sets generated by varying the fitted mechanistic parameters of Healthy Volunteer 1 from ^1^ to 2 times. This restricted perturbation strategy was used to explore nearby dynamical regimes while avoiding parameter combinations that were arbitrarily distant from a parameterization obtained from human data.

The stress-response simulations suggested that, although the model was homeostatically robust over a broad range of conditions, qualitative transitions could occur for particular combinations of stress and modulatory parameters. We therefore performed a formal bifurcation analysis with respect to the stress input *S* and the phenomenological modulatory factors *E*_1_ and *E*_2_.

For this analysis, the explicitly time-dependent model was reduced to an autonomous system.

The circadian CRH drive *T*_1_(*t*) was replaced by its 24-hour mean value,

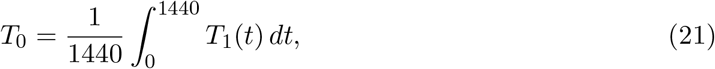

while the periodic modulators *E*_1_(*t*) and *E*_2_(*t*) were replaced by constant values *E*_1_ and *E*_2_.

Sustained stress was represented by a constant input *S*, giving the autonomous CRH equation

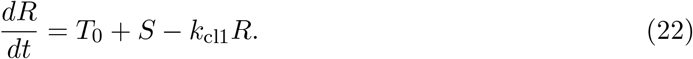

The remaining equations retained the same mechanistic structure and fitted parameters as the full five-state model. Values of *E*_1_ and *E*_2_ were expressed relative to their fitted mean values, *Ē*_1_ and *Ē*_2_, and were varied within the same admissible ranges used in the original model.

Equilibrium branches were continued numerically with *S* as the primary continuation parameter. At each point along the branch, the characteristic roots of the linearized delay differential equation system were calculated and the number of roots with positive real part was recorded. Potential Hopf bifurcations were identified when a complex-conjugate pair of characteristic roots approached and crossed the imaginary axis. Candidate crossings were subsequently corrected numerically to Hopf points using DDE-BifTool. A Hopf crossing was considered confirmed when the corrected point possessed a non-zero oscillation frequency, the real part of the corresponding conjugate eigenvalue pair changed sign across the critical parameter value, and the number of unstable characteristic roots changed by two. Transversality was assessed numerically from the change in the real part of the critical eigenvalues across the bifurcation.

To determine how the stress threshold depended on the modulatory state of the system, corrected Hopf points were subsequently continued in two-parameter space. Hopf loci were computed in the (*S, E*_1_) plane for several fixed values of *E*_2_, and in the (*S, E*_2_) plane for a representative fixed value of *E*_1_. These continuations define sections of a critical surface

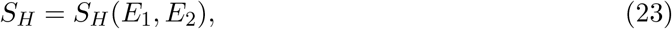

where *S*_*H*_ denotes the sustained-stress level at which equilibrium stability is lost through Hopf bifurcation.

Finally, periodic solutions were initialized directly from selected corrected Hopf points and continued with respect to *S*. Their oscillation amplitude and period were recorded, and their local stability was evaluated using non-trivial Floquet multipliers. Periodic solutions were classified as locally stable when all non-trivial Floquet multipliers remained inside the unit circle. This analysis provided the critical stress thresholds, Hopf bifurcation curves, oscillatory branches, and stability information reported in the Results section.

## Data

All data were obtained from Upton et al [18].

## Author contributions

I.G. Violaris developed the mathematical model, performed the mathematical analysis and proofs, wrote the simulation and fitting code, conducted the computational experiments, analysed the results, and drafted the manuscript.

D. Pinotsis contributed to the interpretation of the modelling results, provided critical feedback on the mathematical and conceptual framework, and contributed to manuscript revision.

M. Tsipouras and K. Kalafatakis contributed equally to the study. They contributed to the study conception and design, interpretation of the biological and modelling results, supervision of the work, and critical revision of the manuscript.

All authors reviewed and approved the final manuscript.

## Supplementary Material

## Appendix A

We consider the reduced three-variable system

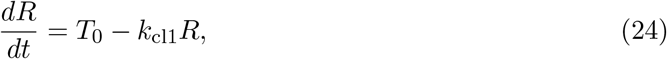

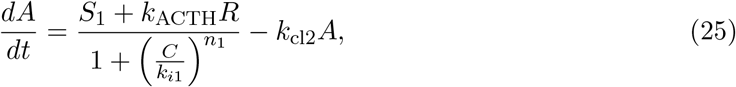

and

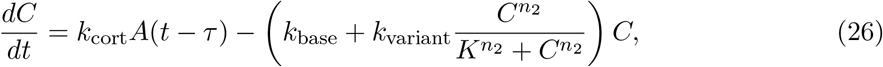

where the CRH driver has been fixed at the constant value *T*_0_ *>* 0.

The first equation can be solved explicitly:

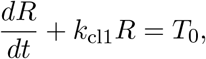

and therefore

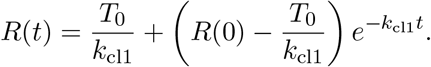

Hence

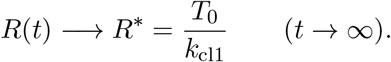

Thus the CRH equation contributes only the stable eigenvalue −*k*_cl1_ and cannot itself gen-erate an oscillatory instability.

Let

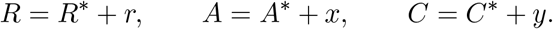

Define

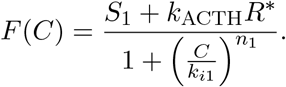

Then

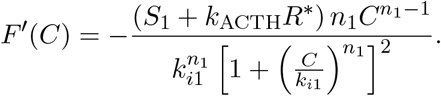

We therefore define

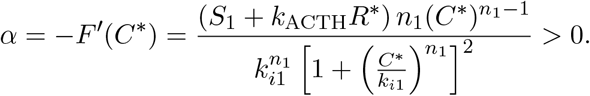

The derivative of the ACTH production term with respect to *R* at equilibrium is

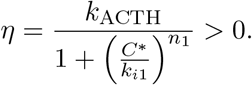

Consequently, the linearized ACTH equation is

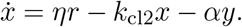

For the cortisol equation, define the complete cortisol-removal function

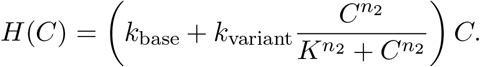

Its derivative is

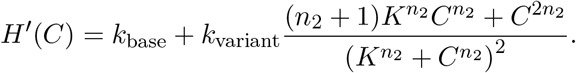

We therefore define

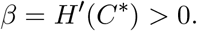

At equilibrium,

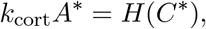

and hence the linearized cortisol equation becomes

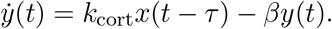

The complete linearized system is therefore

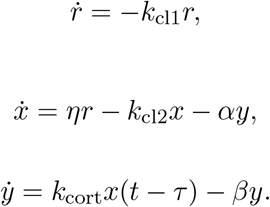

Its characteristic equation factorizes as

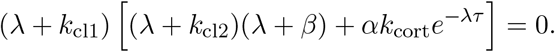

Since *λ* = −*k*_cl1_ *<* 0, any oscillatory instability must arise from

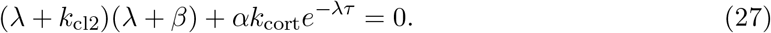

Let

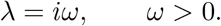

Separating real and imaginary parts gives

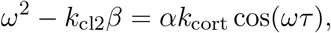

and

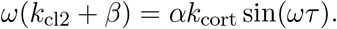

Squaring and adding yields

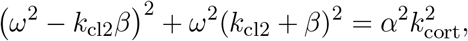

and therefore

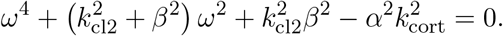

Letting *z* = *ω*^2^ gives

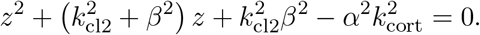

Hence

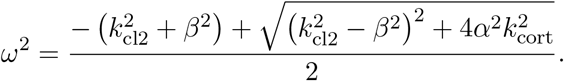

A non-zero real Hopf frequency therefore requires

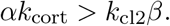

The corresponding critical oscillation period is

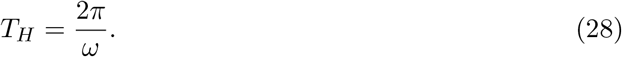

The admissible delay values can be obtained most transparently from the phase of the characteristic equation. Since

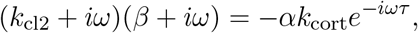

we obtain

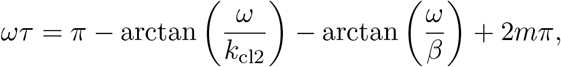

and therefore

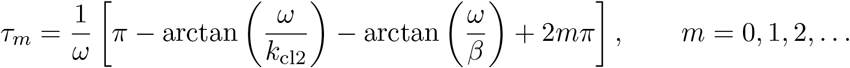

We now consider the relative phase between ACTH and cortisol. At a Hopf point, the first row of the corresponding eigenvector equation is

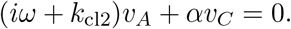

Thus

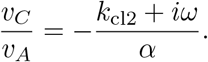

Since *α >* 0, its phase is

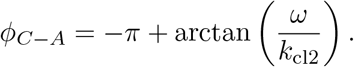

Writing

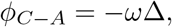

whereΔ denotes the cortisol lag relative to ACTH within one oscillatory period, gives

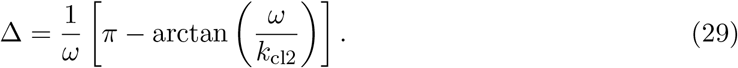

Since

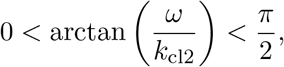

it follows immediately that

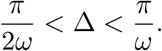

Using *T*_*H*_ = 2*π/ω*, this becomes

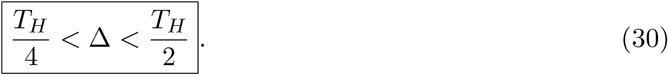

Thus the linearized minimal model imposes an intrinsic lower bound of one quarter of an oscillatory period on the ACTH–cortisol phase lag at a Hopf instability.

For an ultradian period of approximately

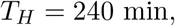

this gives

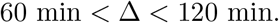

Consequently, an ACTH–cortisol lag of approximately 20 min cannot be reproduced by the Hopf-generated oscillations of this minimal architecture.

## Appendix B

We next consider the five-state model

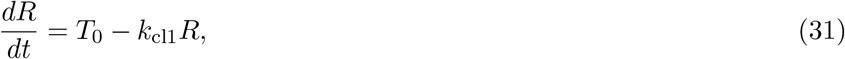

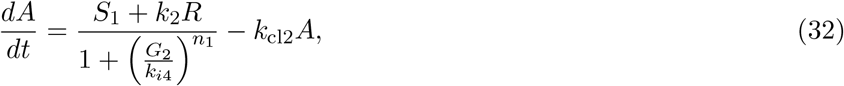

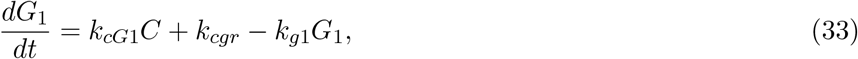

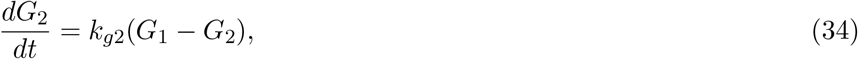

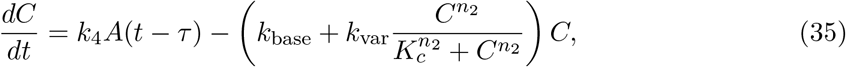

where, as in the analysis of the reduced model, the CRH driver is fixed at a constant value *T*_0_ *>* 0 in order to study the autonomous ultradian dynamics.

The CRH equation is independent of the remaining state variables and has the solution

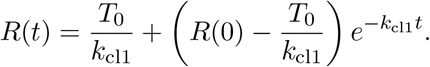

Consequently,

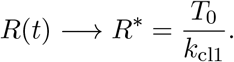

At equilibrium, the remaining state variables satisfy

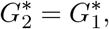

and

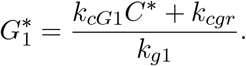

Furthermore,

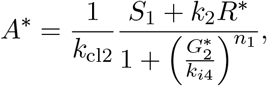

while

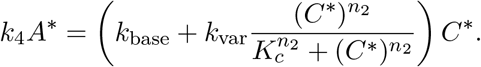

We introduce perturbations

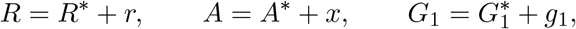

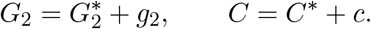

Define the ACTH production function

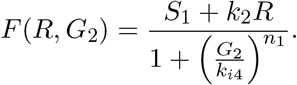

Its derivative with respect to *G*_2_ is

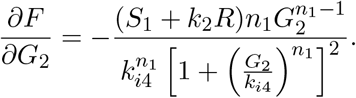

We therefore define

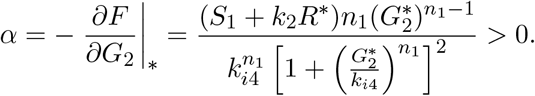

Similarly, define

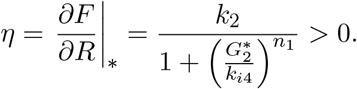

The linearized ACTH equation is then

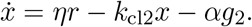

The two intermediate feedback equations are already linear, giving

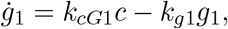

and

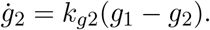

For cortisol, define the complete clearance function

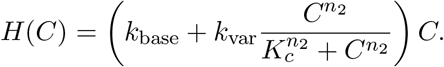

Its derivative is

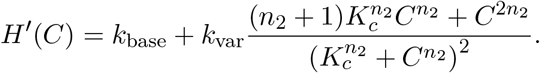

Set

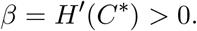

The linearized cortisol equation is therefore

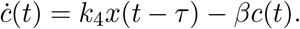

Hence the complete linearized system is

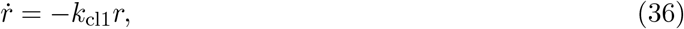

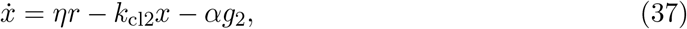

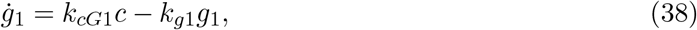

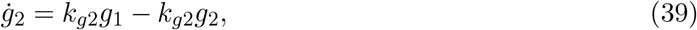

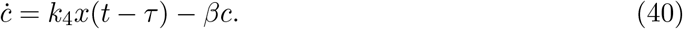

Using the ordering

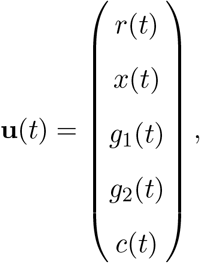

the instantaneous and delayed Jacobian matrices are respectively

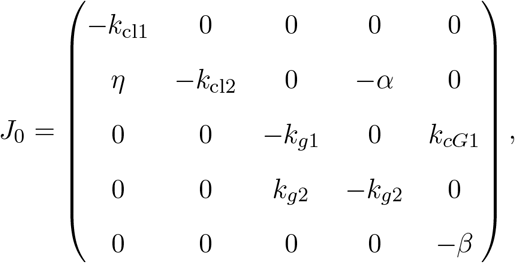

and

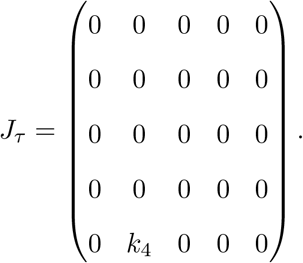

Thus the linearized DDE may be written as

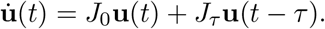

We now seek exponential solutions of the form

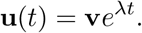

Substitution gives

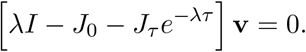

Since the equation for *r* is dynamically decoupled from the feedback loop, one eigenvalue is immediately

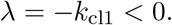

Consequently, the CRH state cannot itself be responsible for an oscillatory instability.

For the remaining four variables, the eigenvalue equations are

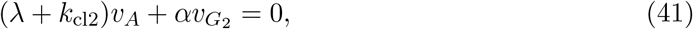

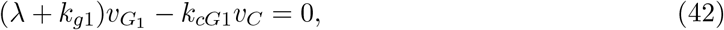

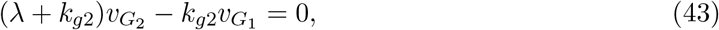

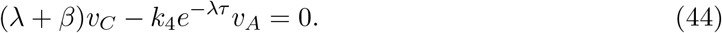

From the cortisol equation,

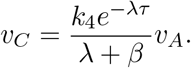

Similarly,

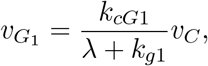

and

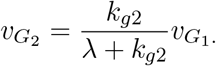

Therefore,

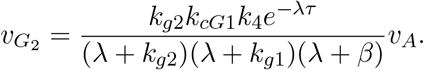

Substituting this expression into the ACTH eigenvalue equation gives

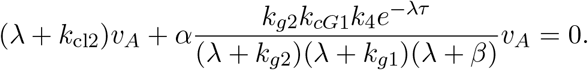

Since *v*_*A*_ ≠ 0 for the oscillatory mode, multiplication by the denominator gives the characteristic equation

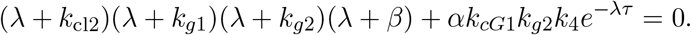

Thus, including the stable CRH eigenvalue, the complete characteristic equation is

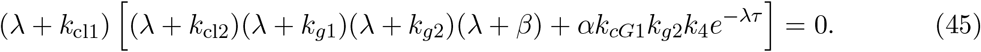

For compactness, define the effective feedback gain

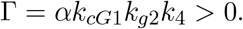

The characteristic equation responsible for oscillatory instability then becomes

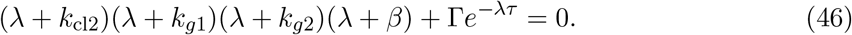

To determine the conditions for a Hopf instability, let

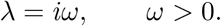

We obtain

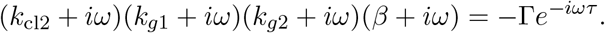

Taking the modulus of both sides gives

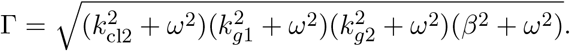

Equivalently,

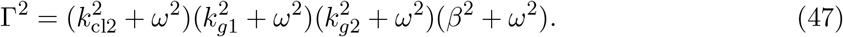

The argument of the left-hand side is

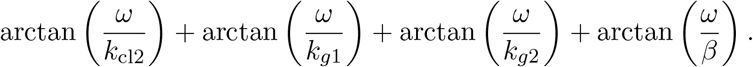

The argument of the right-hand side is

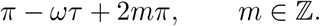

Hence the phase condition for a pair of purely imaginary characteristic roots is

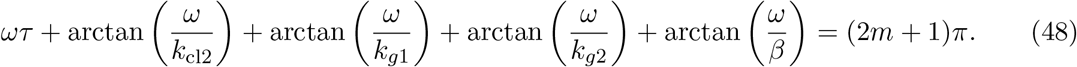

For the principal Hopf branch, *m* = 0, giving

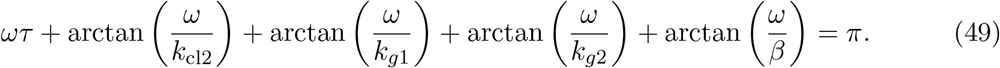

This expression demonstrates the principal architectural difference between the present system and the three-state system. The phase required for closure of the negative-feedback loop is now distributed between the explicit delay and four dynamical stages. In particular, the intermediate variables *G*_1_ and *G*_2_ contribute the additional phase terms

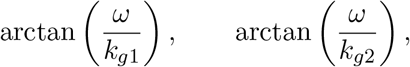

and therefore the required feedback phase need not be supplied by the ACTH–cortisol relationship alone.

We next calculate explicitly the phase difference between ACTH and cortisol. From the cortisol eigenvector equation,

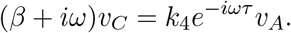

Therefore,

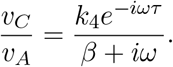

Since *k*_4_ *>* 0, the phase of this ratio is

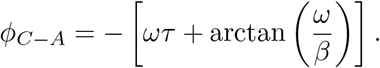

IfΔ_*C*−*A*_ denotes the time by which cortisol lags ACTH on the principal oscillatory branch, then

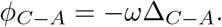

Consequently,

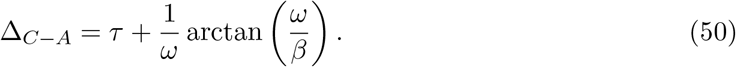

Since

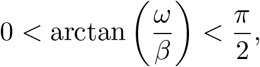

we obtain

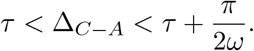

Since the period of the Hopf oscillation is

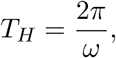

this becomes

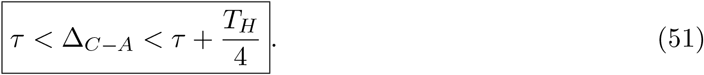

This is fundamentally different from the corresponding result for the three-state architecture, for which the Hopf mode necessarily satisfied

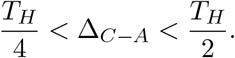

The present architecture therefore removes the structural lower bound of one quarter of an oscillatory period on the ACTH–cortisol lag.

We can demonstrate this explicitly for the physiologically relevant case

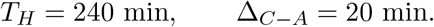

The required angular frequency is

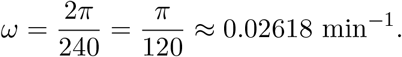

For example, take an explicit ACTH–cortisol delay of

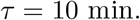

The desired total lag of 20 min requires

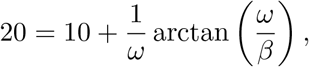

so that

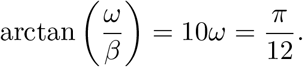

Consequently,

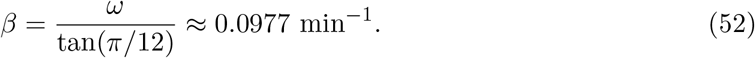

For these values,

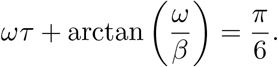

The remaining three dynamical stages must therefore contribute

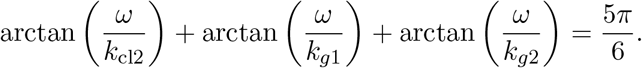

To give a simple constructive example, let

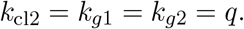

Then

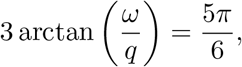

and therefore

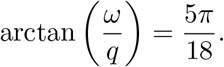

Thus

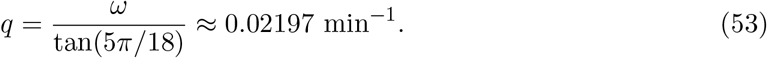

It follows that the positive parameter values

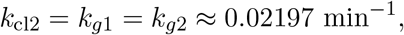

together with

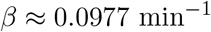

and *τ* = 10 min satisfy exactly the phase condition for a Hopf oscillation with period 240 min and an ACTH–cortisol lag of 20 min.

The corresponding magnitude condition determines the required critical feedback gain:

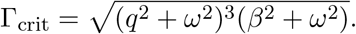

Numerically,

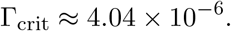

Since

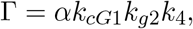

the gain condition can therefore be written as

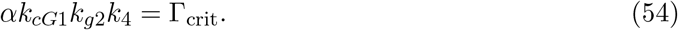

Hence both the phase and magnitude conditions for purely imaginary roots can be satisfied simultaneously while preserving the desired ultradian period and ACTH–cortisol lag.

It remains to verify that the imaginary roots cross the imaginary axis rather than merely touching it. Define

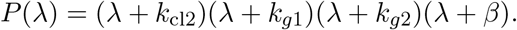

The characteristic equation is

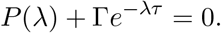

Differentiating implicitly with respect to the effective feedback gain Γ gives

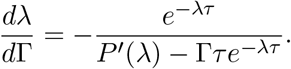

At a characteristic root,

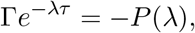

and hence

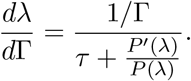

Since

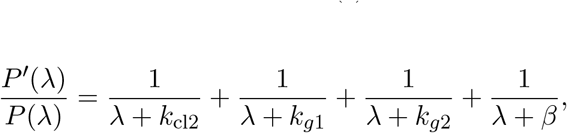

we obtain, at *λ* = *iω*,

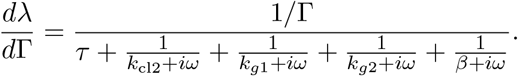

The real part of the denominator is

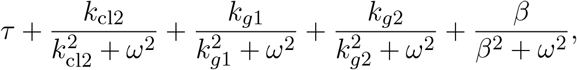

which is strictly positive. Therefore,

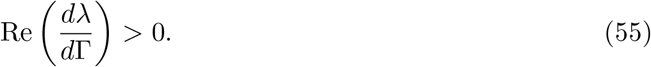

Thus the conjugate pair of characteristic roots crosses the imaginary axis transversely as the effective feedback gain passes through its critical value.

Furthermore, for *ω >* 0 the magnitude function

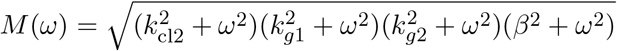

is strictly increasing in *ω*. Hence, for a fixed positive critical gain Γ, the magnitude condition admits at most one positive Hopf frequency. The pair *λ* = ±*iω* is therefore isolated.

We conclude that, unlike the three-state system, the five-state feedback architecture admits a Hopf instability having simultaneously an ultradian period of approximately 240 min and an ACTH–cortisol phase lag of approximately 20 min. The additional *G*_1_ and *G*_2_ states provide the phase accumulation required to close the negative feedback loop without forcing the cortisol oscillation to lag ACTH by at least one quarter of a cycle. Thus the physiological ACTH– cortisol lag and the observed ultradian timescale are dynamically compatible within the five-state architecture.

## Appendix C

In this section we present the fits produced by the model for 3 more eligible volunteers from Upton et al [18]

## Appendix D

In this section we present the fits of the modulated model for Healthy Volunteers 4–6.

## References

[1] Russell, G. M., Kalafatakis, K., & Lightman, S. L. (2015). The importance of biological oscillators for hypothalamic-pituitary-adrenal activity and tissue glucocorticoid response: coordinating stress and neurobehavioural adaptation. Journal of neuroendocrinology, 27(6), 378–388. 10.1111/jne.12247

[2] Weitzman, E. D., Fukushima, D., Nogeire, C., Roffwarg, H., Gallagher, T. F., & Hellman, L. (1971). Twenty-four hour pattern of the episodic secretion of cortisol in normal subjects. The Journal of clinical endocrinology and metabolism, 33(1), 14–22. 10.1210/jcem-33-1-14

[3] Lightman, S. L., & Conway-Campbell, B. L. (2010). The crucial role of pulsatile activity of the HPA axis for continuous dynamic equilibration. Nature reviews. Neuroscience, 11(10), 710–718. 10.1038/nrn2914

[4] Kadmiel, M., & Cidlowski, J. A. (2013). Glucocorticoid receptor signaling in health and disease. Trends in pharmacological sciences, 34(9), 518–530. 10.1016/j.tips.2013.07.003

[5] Sapolsky, R. M., Romero, L. M., & Munck, A. U. (2000). How do glucocorticoids influence stress responses? Integrating permissive, suppressive, stimulatory, and preparative actions. Endocrine reviews, 21(1), 55–89. 10.1210/edrv.21.1.0389

[6] Gupta, S., Aslakson, E., Gurbaxani, B. M., & Vernon, S. D. (2007). Inclusion of the glucocorticoid receptor in a hypothalamic pituitary adrenal axis model reveals bistability. Theoretical biology & medical modelling, 4, 8. 10.1186/1742-4682-4-8

[7] Walker, J. J., Terry, J. R., & Lightman, S. L. (2010). Origin of ultradian pulsatility in the hypothalamic-pituitary-adrenal axis. Proceedings. Biological sciences, 277(1688), 1627–1633. 10.1098/rspb.2009.2148

[8] Vinther, F., Andersen, M., & Ottesen, J. T. (2011). The minimal model of the hypothalamic-pituitary-adrenal axis. Journal of mathematical biology, 63(4), 663–690. 10.1007/s00285-010-0384-2

[9] Kim, L. U., D’Orsogna, M. R., & Chou, T. (2018). Perturbing the Hypothalamic-Pituitary-Adrenal Axis: A Mathematical Model for Interpreting PTSD Assessment Tests. Computational psychiatry (Cambridge, Mass.), 2, 28–49. 10.1162/CPSYa00013

[10] Herman, J. P., McKlveen, J. M., Ghosal, S., Kopp, B., Wulsin, A., Makinson, R., Scheimann, J., & Myers, B. (2016). Regulation of the Hypothalamic-Pituitary-Adrenocortical Stress Response. Comprehensive physiology, 6(2), 603–621. 10.1002/cphy.c150015

[11] Keenan, D. M., Licinio, J., & Veldhuis, J. D. (2001). A feedback-controlled ensemble model of the stress-responsive hypothalamo-pituitary-adrenal axis. Proceedings of the National Academy of Sciences of the United States of America, 98(7), 4028–4033. 10.1073/pnas.051624198

[12] Regazzo, D., Mondin, A., Scaroni, C., Occhi, G., & Barbot, M. (2022). The Role of Glucocorticoid Receptor in the Pathophysiology of Pituitary Corticotroph Adenomas. International journal of molecular sciences, 23(12), 6469. 10.3390/ijms23126469

[13] Charmandari, E., Chrousos, G. P., Lambrou, G. I., Pavlaki, A., Koide, H., Ng, S. S., & Kino, T. (2011). Peripheral CLOCK regulates target-tissue glucocorticoid receptor transcriptional activity in a circadian fashion in man. PloS one, 6(9), e25612. 10.1371/journal.pone.0025612

[14] Miller, W. L., & Auchus, R. J. (2011). The molecular biology, biochemistry, and physiology of human steroidogenesis and its disorders. Endocrine reviews, 32(1), 81–151. 10.1210/er.2010-0013

[15] van der Spoel, E., Roelfsema, F., Akintola, A. A., Jansen, S. W., Slagboom, P. E., Westendorp, R. G. J., Blauw, G. J., Pijl, H., & van Heemst, D. (2020). Interrelationships Between Pituitary Hormones as Assessed From 24-hour Serum Concentrations in Healthy Older Subjects. The Journal of clinical endocrinology and metabolism, 105(4), e1201–e1214. 10.1210/clinem/dgz253

[16] Violaris, I. G., Kalafatakis, K., Zavala, E., Tsoulos, I. G., Lampros, T., Lightman, S. L., Tsipouras, M. G., Giannakeas, N., Tzallas, A., & Russell, G. M. (2021). Modelling Hydrocortisone Pharmacokinetics on a Subcutaneous Pulsatile Infusion Replacement Strategy in Patients with Adrenocortical Insufficiency. Pharmaceutics, 13(6), 769. 10.3390/pharmaceutics13060769

[17] de Lacerda, L., Kowarski, A., & Migeon, C. J. (1973). Diurnal variation of the metabolic clearance rate of cortisol. Effect on measurement of cortisol production rate. The Journal of clinical endocrinology and metabolism, 36(6), 1043–1049. 10.1210/jcem-36-6-1043

[18] Upton, T. J., Zavala, E., Methlie, P., Kämpe, O., Tsagarakis, S., Øksnes, M., Bensing, S., Vassiliadi, D. A., Grytaas, M. A., Botusan, I. R., Ueland, G., Berinder, K., Simunkova, K., Balomenaki, M., Margaritopoulos, D., Henne, N., Crossley, R., Russell, G., Husebye, E. S., & Lightman, S. L. (2023). High-resolution daily profiles of tissue adrenal steroids by portable automated collection. Science translational medicine, 15(701), eadg8464. 10.1126/scitranslmed.adg8464

[19] Waite, E. J., McKenna, M., Kershaw, Y., Walker, J. J., Cho, K., Piggins, H. D., & Lightman, S. L. (2012). Ultradian corticosterone secretion is maintained in the absence of circadian cues. The European journal of neuroscience, 36(8), 3142–3150. 10.1111/j.1460-9568.2012.08213.x

[20] Jacobson, L., Muglia, L. J., Weninger, S. C., Pacák, K., & Majzoub, J. A. (2000). CRH deficiency impairs but does not block pituitary-adrenal responses to diverse stressors. Neuroendocrinology, 71(2), 79–87. 10.1159/000054524

